# An NKX2-5 homolog is required downstream of BMP signaling to pattern the sensory-adhesive organ of a tunicate larva

**DOI:** 10.64898/2025.12.18.695272

**Authors:** Christopher J. Johnson, Joshua Kavaler, Christina D. Cota, Alberto Stolfi

## Abstract

Tunicates are the sister group to vertebrates within the chordate phylum, yet unlike other chordate groups, they evolved a biphasic lifecycle alternating between motile larvae and sessile adults. The papillae of most tunicate larvae are the key sensory-adhesive organ regulating their settlement and metamorphosis. The papillae are nearly always arranged as a group of three morphologically identical organs that arise from an anterior neural plate border region nested between ventral epidermis and more dorsal/posterior neural tube progenitors. Due to their embryonic origin and molecular signatures, this anterior border has been evolutionarily linked to vertebrate placode regions. It was previously shown that the specification, patterning, and morphogenesis of the embryonic papilla region all depend on BMP signaling, though downstream mechanisms remain poorly understood. Here we show that the NKX2-3/5/6 ortholog NK4 is a key transcription factor that acts downstream of BMP signaling to pattern the papillae in the *Ciona* embryo. We present evidence that *NK4* is activated by BMP signaling and encodes a transcriptional repressor that is required to restrict the expression of the papilla regulatory gene *Foxg* to three cell clusters that give rise to the three papillae. Loss of NK4 function results in the formation of a single large papilla. In contrast, overexpression of NK4 represses *Foxg*, eliminating the papillae. We also show that the expression of *NK4* is restricted dorsally by the BMP antagonist Chordin, while the ventrally-expressed transcription factor Msx alleviates the repressive effect of NK4, potentially allowing for the characteristic tripartite patterning of the papillae.

## INTRODUCTION

Tunicates have emerged as model organisms for the study of chordate evolution and development (Anselmi et al., 2025). Despite sharing key anatomical, cellular, and molecular traits with other chordate subphyla, a majority of tunicate species alternate between motile, pelagic larval and sessile, benthic adult lifecycle phases (Karaiskou et al., 2015; Lemaire, 2011). The sensory-adhesive papilla organ of the larva plays an indispensable role in driving substrate attachment and metamorphosis in most species (Nakayama-Ishimura et al., 2009). In solitary species studied in the lab, it has been shown that continuous attachment to a solid substrate is necessary and sufficient to trigger metamorphosis (Matsunobu and Sasakura, 2015). This metamorphosis has been shown to be driven by the mechanosensory abilities of the papillae, and direct mechanical or optogenetic stimulation of the papilla neurons (PNs) can substitute for attachment in initiating metamorphosis (Totsuka and Hotta, 2025; Wakai et al., 2021). It is thought that the PNs sense mechanical forces generated and/or augmented by substrate attachment (Hozumi et al., 2025), which in turn depends on adhesive material-secreting collocytes that are present alongside the PNs in the papilla organ (Zeng et al., 2019a; Zeng et al., 2019b). Consistent with this model, metamorphosis is blocked by genetically eliminating PNs or perturbing their function (Sakamoto et al., 2022). In most species, the papilla organ is invariably organized in a tripartite manner, consisting of three constituent papillae arranged as vertices of a triangle: two dorsal, lateral papillae and one ventral, medial papilla (Berrill, 1930; Dolcemascolo et al., 2009; Turon, 1991). Each papilla appears to consist of the same number and arrangement of cell types, of which at least five are known based on cytological and molecular characterization (Johnson et al., 2024; Zeng et al., 2019b). This tripartite arrangement is highly conserved, even in the much larger larvae of certain compound tunicate species (Cloney, 1977), suggesting some adaptive advantage for the larva.

The papillae arise from the anteriormost cells of the neural plate, forming an anterior neural border (ANB) that might be evolutionarily connected to the anterior cranial placodes of vertebrates (Abitua et al., 2015; Feinberg et al., 2019; Ikeda et al., 2013; Liu et al., 2023; Roure et al., 2023; Roure et al., 2025; Thawani and Groves, 2020; Tresser et al., 2010; Wagner and Levine, 2012). In multiple species studied, the specification and patterning of the papilla region is regulated by BMP signaling (Liu et al., 2023; Roure et al., 2023). First, BMP signaling is required for proper papilla specification, though in more anterior/ventral epidermis it may suppress papilla fate (Liu et al., 2023; Roure et al., 2023). Later, BMP signaling patterns the papilla territory into its familiar tripartite arrangement (Liu et al., 2023; Roure et al., 2023). It has also been shown that the tripartite organization of the papillae depends on the precise specification of three discrete, Foxg+/Islet+ cell clusters at the mid/late tailbud stages (Johnson et al., 2024; Liu and Satou, 2019; Roure et al., 2023; Wagner et al., 2014). However, our understanding of how BMP or other candidate pathways accomplish this remains incomplete (**Figure 1**).

**Figure 1.**
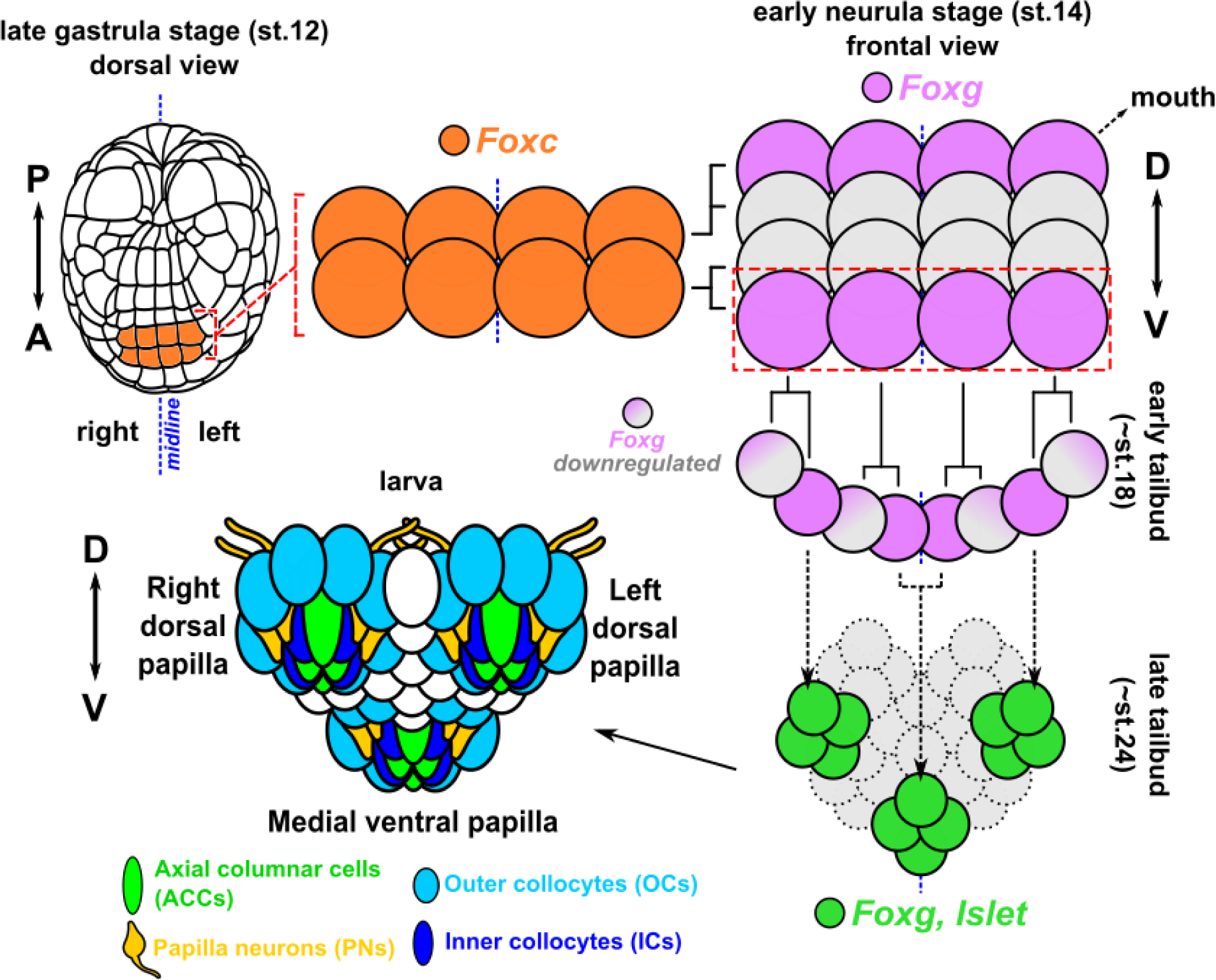
Summary diagram of papilla development in *Ciona.* A stylized diagram summarizing what is currently know about the cell lineages and major cell fate specification events in the developing papilla territory of *Ciona* embryos. Briefly, the papillae are derived from anterior-most *Foxc*-expressing cells of the anterior neural plate border at gastrula stage. The anterior-most row of daughter cells (red dashed rectangle) go on to express *Foxg* at the early neurula stage. During tailbud stages, this row of cells begins to bend ventrally, forming a “U”-shaped swatch of cells. *Foxg* expression is maintained in three discrete clusters of cells and downregulated in the remaining cells of this row. Eventually, sustained Foxg activates *Islet* in the three cell clusters and their descendants, giving rise to the three protruding sensory-adhesive papillae of the hatched larva: one medial, one ventral papilla, and two lateral, dorsal papillae. Each papilla is composed of invariant numbers of different cell types (4 axial columnar cells, 4 inner collocytes, 4 papilla neurons, and 8 outer collocytes, for a total of 20 cells). See text for references.

Here we show that the BMP-responsive regulatory gene *Nkx2-3/5/6*, the ortholog of *NKX2-5, NKX2-3*, and *NKX2-6* in humans, and of *tinman* in *Drosophila*, is a key intermediate regulator that mediates the BMP-dependent tripartite patterning of the *Ciona* papillae. We show that BMP signaling in the papilla territory promotes expression of *Nkx2-3/5/6* (which we call *NK4* herein, based on the original name of the *Drosophila* tinman protein, NK-4)(Kim and Nirenberg, 1989). We further show that NK4 acts as a transcriptional repressor that is both necessary and sufficient to downregulate *Foxg* expression in specific sub-regions of this territory, thereby allowing for the formation of discrete *Foxg+* cell clusters that give rise to the three papillae. Disrupting *NK4* largely phenocopies BMP inhibition, resulting in a single large papilla. We discuss our findings in the context of a larger model that we propose as underlying the development of the conserved tripartite organization of the papilla organ, and its implications for larval settlement and metamorphosis.

## RESULTS

### Knocking out NK4 in the papillae using CRISPR/Cas9 phenocopies BMP inhibition

We initially identified NK4 as a potential regulator of cell fate in the papillae. *NK4* is expressed throughout the papilla territory (**Figure 2A, Figure S1, Video S1**)(Johnson et al., 2020), where it was hypothesized to be important for the specification of the Axial Columnar Cells (ACCs), a papilla cell type with sensory capabilities but of unknown function (Hoyer et al., 2024; Johnson et al., 2020). We thus originally sought to knock out *NK4* specifically in the papilla territory, to assay its role in ACC specification. To do so, we selected a previously designed and validated, high-efficiency single-chain guide RNA (sgRNA) targeting the second exon of *NK4* (**Figure 2B**), predicted to cut within the sequence encoding part the DNA-binding homeodomain (Gandhi et al., 2017). To limit CRISPR/Cas9 activity to the early papilla cell lineages, we used the *Foxc* promoter to drive expression of Cas9 (*Foxc>Cas9*)(Johnson et al., 2024). When we co-electroporated the sgRNA expression plasmid (*U6>NK4.3*) with *Foxc>Cas9*, we unexpectedly observed a “*Cyrano*” phenotype (Roure et al., 2023) in which a single protruding papilla forms in place of the familiar three papillae arranged as a triangle (**Figure 2C,D**). This phenotype was originally observed in larvae subjected to BMP signaling knockdown (Liu et al., 2023; Roure et al., 2023), but to our knowledge this is the first report of a CRISPR condition with the same or similar phenotype. The ACC reporter gene *CryBG>GFP* was still expressed in *NK4* CRISPR larvae showing a clear single-papilla phenotype (**Figure 2C,E**), suggesting that NK4 is specifically involved in papilla organ patterning but not necessarily cell type specification or differentiation. To test this, we replicated the CRISPR-mediated disruption of *NK4* while assaying the expression of reporters that label the other cell types of the papillae (**Figure 3A,B**). Indeed, all cell type reporters were still labeling cells contributing to the resulting single large papilla. The same phenotype of a single large papilla was obtained using another sgRNA targeting a different part of the *NK4* gene, further supporting our observations (**Figure S2**). Despite only having one large papilla, *NK4* CRISPR larvae were still fully capable of undergoing metamorphosis (**Figure 3C,D**), likely due to the fact that all the cell types, especially the PNs, were still present. Of note, PNs in *NK4 CRISPR* larvae still had their apical cilia, which are hypothesized to be important for mechanosensory ability (**Video S2-4**) These results suggest that the tripartite organization of the papillae might not be required for metamorphosis under laboratory conditions, but could be more important for robust adhesion or to fine-tune substrate selection in the wild.

**Figure 2.**
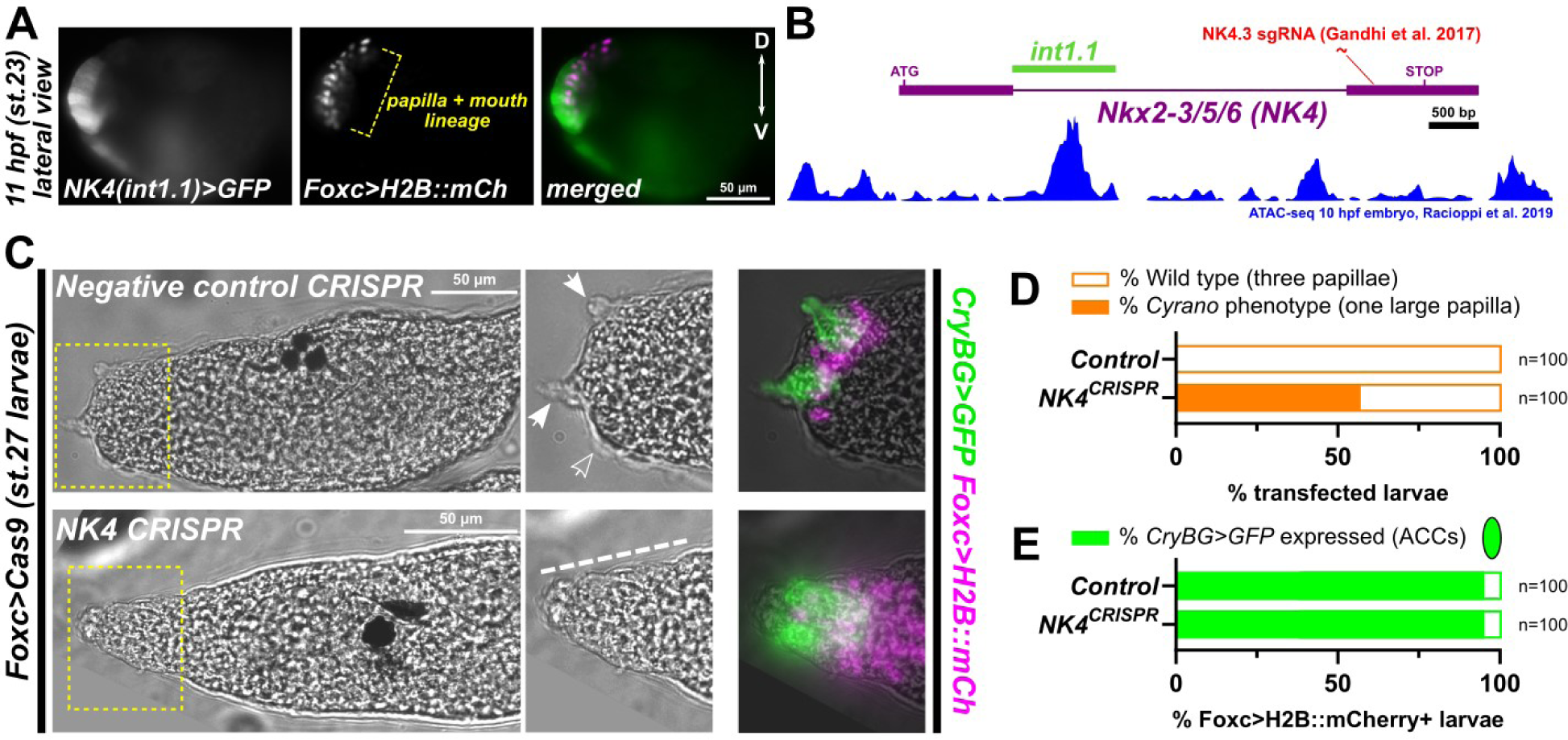
*NK4*, an *NKX2-3/5/6* ortholog, is required for the tripartite patterning of the papillae. A) A GFP reporter plasmid constructed using a portion of the sole intron of the *NK4* gene of *Ciona robusta* (“int1.1”), fused to the basal promoter of the *FOG* gene (Rothbächer et al., 2007). See **Supplemental Sequences** file for details), is expressed throughout the Foxc+ papilla territory in stage 23 embryos, recapitulating the previously reported mRNA *in situ* hybridization pattern. Int1.1 reporter activity can also be seen in ventral epidermis and trunk endoderm. hpf = hours post-fertilization. B) Diagram depicting the *NK4* locus and location of both the int1.1 *cis*-regulatory element and the target of the published *NK4.3* sgRNA. Underneath is the ATAC-seq profile of 10 hpf embryos, from Racioppi et al. 2019, showing a peak of DNA accessibility centered on the int1.1 element. C) Papilla lineage-specific CRISPR/Cas9-mediated mutagenesis of *NK4* showing the single, enlarged papilla protrusion (indicated by dashed line) similar to the *Cyrano* phenotype described in Roure et al. 2023 and replicated in Liu et al. 2023. Compare to the negative control CRISPR using the non-targeting *“Control”* sgRNA instead. Right panels: the *CryBG>GFP* reporter (green) reveals the presence of ACCs in the single papilla of the *NK4* CRISPR larvae. Left panels show brightfield images (zoomed in panel indicated in yellow dashed outline), while right panels show brightfield overlaid on GFP/mCherry fluorescence. D) Scoring the percentage of transfected larvae showing the single large papilla phenotype (“*Cyrano*”) in *NK4* CRISPR vs. negative control larvae electroporated with only the reporter plasmids. Transfection was monitored using a combination of *Islet>GFP* and *Foxc>H2B::mCherry* reporter plasmid expression in the papillae. E) Scoring the proportion of transfected (*Foxc>H2B::mCherry+*) larvae showing *CryBG>GFP* expression as shown in panel C. No difference was observed in the proportion of larvae expressing *CryBG>GFP* between *NK4* CRISPR and control larvae. All GFP sequences are tagged with Unc-76, which promotes cytoplasmic localization over nuclear localization (Dynes and Ngai, 1998). Scale bars = 50 m. mCh: mCherry; D: dorsal; V: ventral.

**Figure 3.**
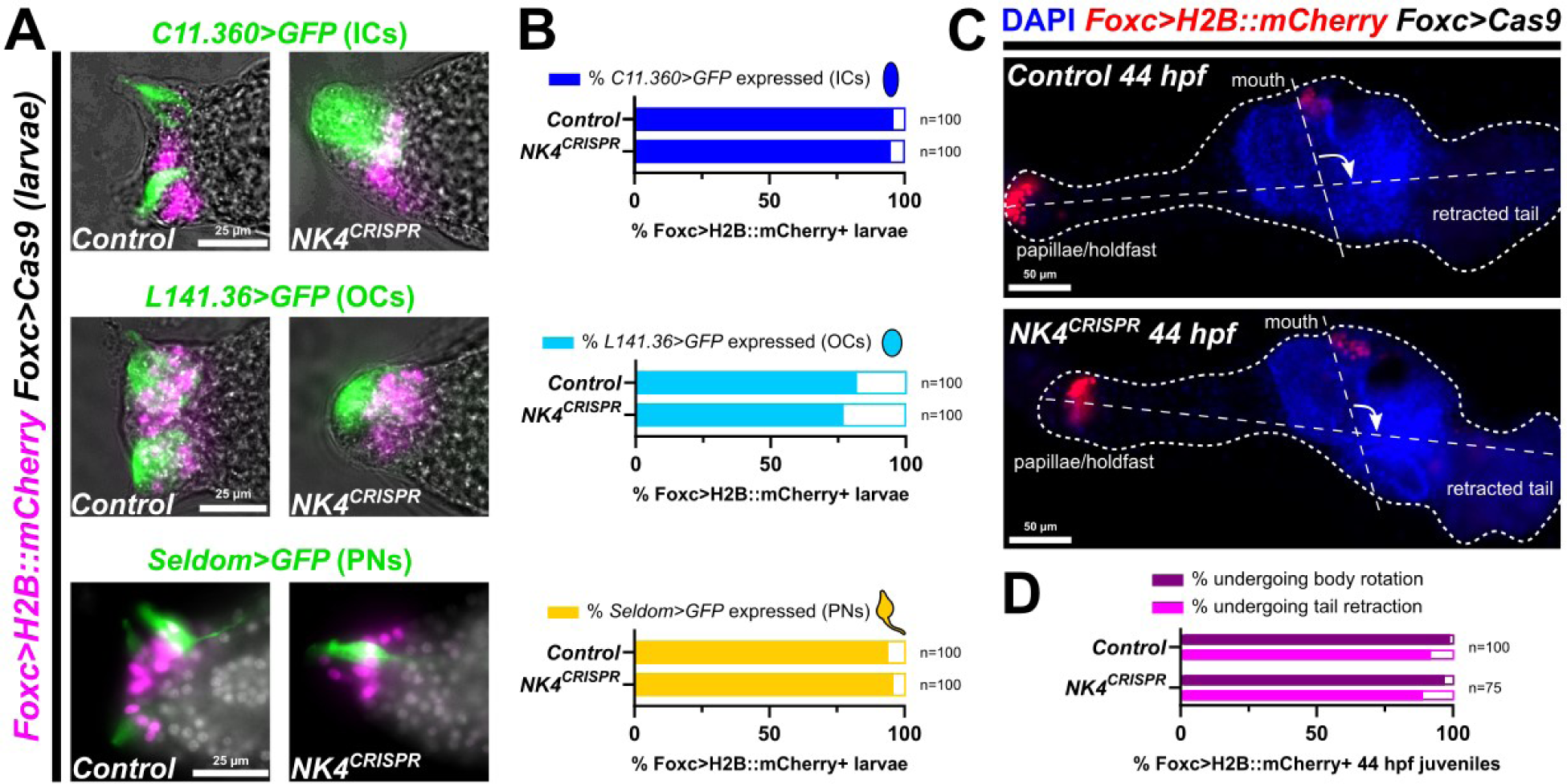
*NK4* CRISPR does not significantly impact specification of other papilla cell types or metamorphosis, despite resulting in a single large papilla. A) Papilla lineage-specific *CRISPR* mutagenesis of *NK4* does not result in loss of other papilla cell type-specific reporter gene expression in spite of the loss of the typical tripartite organization of the papillae. ICs: inner collocytes; OCs: outer collocytes; PNs: papilla neurons. Counterstain in *Seldom>GFP* images is DAPI (white overlay). Scale bars = 25 µm. IC and PN images acquired at 17 hours post-fertilization (hpf; St. 27), while OC images were acquired at 21 hpf (St. 29) on account of the late onset of *L141.36* expression as previously reported (Johnson et al., 2024). B) Scoring the percentage of transfected larvae showing expression of each cell type-specific reporter, comparing *NK4* CRISPR vs. negative control larvae as indicated in panel A. C) *NK4* CRISPR in the papilla lineage also does not significantly prevent metamorphosis, as assayed in 44 hpf juveniles (St. 35) undergoing tail retraction and body axis rotation (white arrow). Scale bars = 50 µm. D) Scoring the percentage of transfected juveniles undergoing either metamorphic process as depicted in panel C. All GFP sequences tagged with Unc-76.

### BMP signaling activates *NK4* expression in the papilla territory

In *Ciona, NK4* has been shown to be activated by BMP signaling emanating from ventral regions (Christiaen et al., 2010; Roure et al., 2023; Waki et al., 2015). In the migrating cardiopharyngeal progenitors, BMP signaling activates *NK4*, which in turn modulates cell migration and fate decisions in this lineage (Christiaen et al., 2010). In other studies, *NK4* expression in the ventral epidermis was shown to be upregulated by BMP treatment and downregulated by pharmacological inhibitors of the BMP pathway (Roure et al., 2023; Waki et al., 2015). Because the characteristic phenotype originally described in BMP-inhibited larvae so closely resembled that of our *NK4* CRISPR larvae, we reasoned that *NK4* might act downstream of BMP signaling to pattern the papillae.

To test this, first we showed that overexpression of the BMP antagonist Noggin (*Foxc>Noggin)* also results in a single protruding papilla containing all known papilla cell types (**Figure 4A**). Second, we found that *Foxc>Noggin* also greatly reduces the expression of the *NK4>GFP* reporter plasmid in the papillae (**Figure 4B**). Expression was not completely abolished, but quantification of mean GFP fluorescence (normalized to whole-larva expression of an *Islet>mCherry* reporter) showed that it is significantly reduced upon BMP inhibition (**Figure 4C-E, Figure S3**). Interestingly, we observed a complete loss of expression of the the BMP-dependent ventral papilla marker gene *Msx* (Roure et al., 2023) in Noggin-overexpression larvae, compared to a more modest reduction in *NK4* CRISPR larvae (**Figure 4F,G**). Despite the presence of a single large papilla in in many of these larvae, *Msx* reporter plasmid expression was still observed in the ventral area of the papilla region (**Figure 4F**). This suggests that, although NK4 functions downstream of BMP signaling to regulate the tripartite organization of the papillae, activation of *Msx* in the ventral papilla by BMP is mostly independent of NK4.

**Figure 4.**
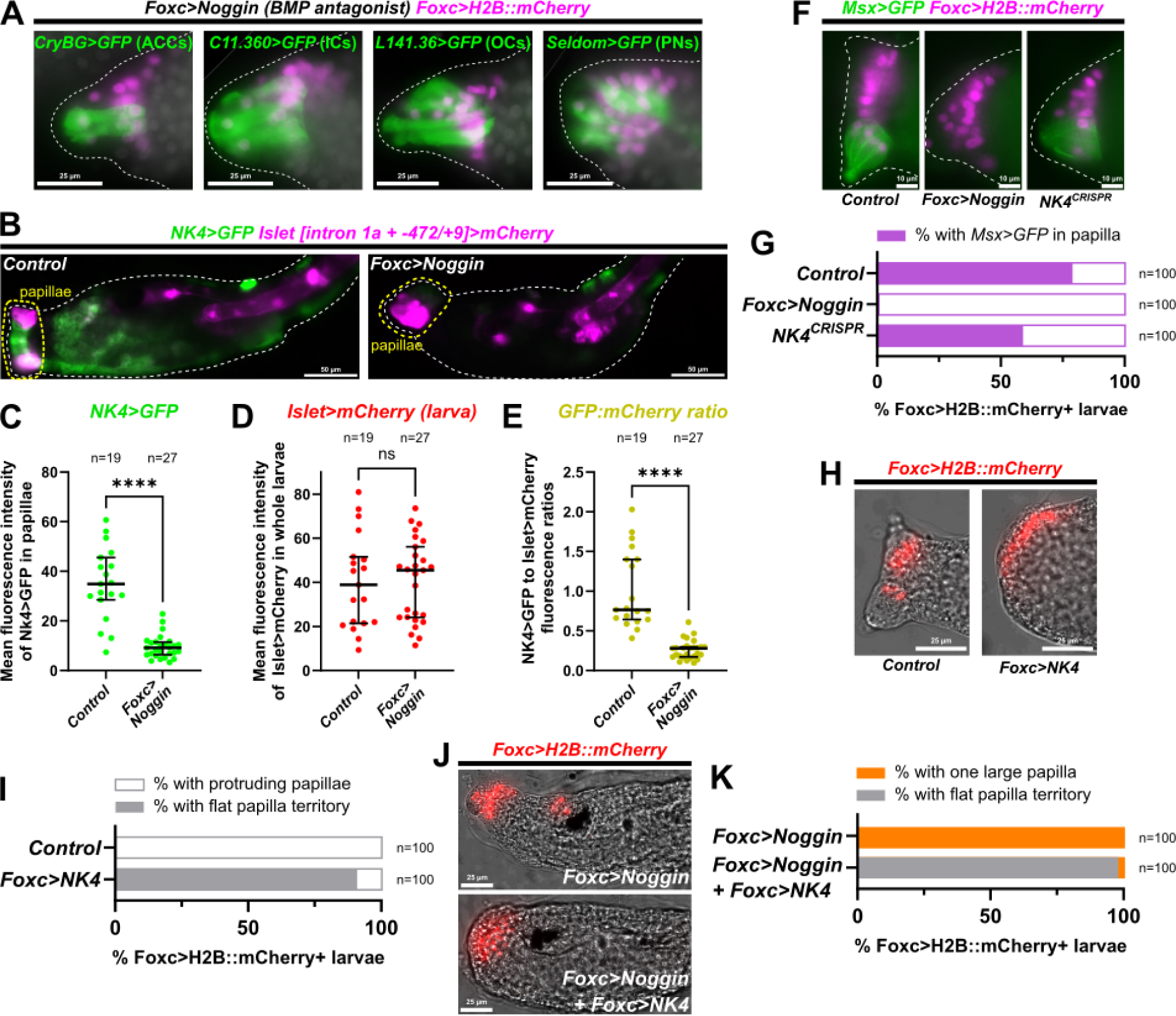
NK4 acts downstream of BMP signaling in the papillae. A) Overexpression of the BMP antagonist Noggin in the papilla lineage (using *Foxc>Noggin* plasmid) results in a “*Cyrano”* phenotype similar to *NK4* CRISPR: a single large papilla containing all four papilla cell types (ACCs, ICs, OCs, adn PNs). DAPI counterstain shown in white overlay. All at 17 hpf (St. 27) except *L141.36* reporter expression, which is visible only around 21 hpf (St. 29). B) *NK4>*GFP reporter expression (green) was quantified by measuring mean fluorescence in the papilla territory (yellow dashed area), in negative control and *Foxc>Noggin-*electroporated larvae. Transfection efficacy was normalized using whole-larva expression of *Islet [intron 1a + −472/+9]>mCherry* (magenta), which is expressed in papilla and notochord cells. Scale bars = 50 µm. C) Comparison of mean fluorescence intensity readouts of *NK4>GFP* expression in the papilla region (as indicated in panel B), between negative control and *Foxc>Noggin* larvae. Data show significant downregulation of *NK4* reporter expression in the papillae by Noggin overexpression. 5 outliers of fluorescence over 80 were removed from the control dataset. **** p<0.0001 by two-tailed Mann-Whitney test. D) Comparison of mean fluorescence intensity readouts of *Islet>mCherry* fluorescence in the whole larva, between the same larvae plotted in panel C and shown in panel B. Data show that overexpression of Noggin had no significant effect on Islet expression levels throughout the larva, indicating comparable transfection efficacies and suitable use of whole-larva *Islet* reporter expression to normalize *NK4* reporter activity. p = 0.7404 by two-tailed Mann-Whitney. E) Ratio of GFP to mCherry intensity values plotted in panels C and D, showing that *NK4* reporter is significantly downregulated by Noggin overexpression even when normalized by transfection efficacy. p<0.0001 by two-tailed Mann-Whitney. Outliers with GFP:mCherry ratio >3.0 (three in the control condition) were removed from the entire dataset. Complete data with flagged outliers can be found in **Supplemental Data S1.** Normalization to papilla-specific *Islet>mCherry* expression can be found in **Figure S3.** F) Expression of the *Msx>CD4::GFP* reporter (green) in negative control CRISPR, Noggin overexpression, and *NK4* CRISPR (using *Foxc>Cas9*). G) Comparison of percentage of transfected larvae showing *Msx* reporter expression in the ventral papilla region, comparing the conditions shown in panel F. H) Overexpression of NK4 using the *Foxc* promoter results in the absence of protruding papillae and flattening of the entire anterior epidermis. I) Scoring the percentage of transfected larvae showing the flattened papilla territory, comparing the conditions in panel H. J) Overexpression of NK4 in the papilla region can override the Noggin overexpression phenotype, converting a single large papilla into a flattened epidermis. All scale bars (except in panel B) = 25 µm. All GFP sequences except *Msx>CD4::GFP* are tagged with Unc-76.

In contrast to *NK4* CRISPR or BMP inhibition, overexpression of NK4 (*Foxc>NK4*) resulted in the near opposite of the *Cyrano* phenotype: a flat layer of cells where the papillae normally form (**Figure 4H,I**). When combining NK4 overexpression with BMP inhibition (Noggin overexpression), the NK4 overexpression phenotype (flat/no papillae) clearly prevailed (**Figure 4J,K**). Taken together, these data suggest that NK4 acts downstream of BMP signaling in the papilla territory to pattern this region into the three protrusions invariably seen in wild-type larvae.

### NK4 represses *Foxg*

Having established a role for NK4 in patterning the papilla organ downstream of BMP signaling, we next sought to better understand exactly how NK4 achieves this. *NK4* expression has been shown to begin during the neurula stage (Imai et al., 2004). By mRNA *in situ* hybridization we observed *NK4* expression in the middle of the papilla territory during the late neurula stage, overlapping with *Foxg* expression when the latter is still expressed in the anteriormost “U”-shaped row of cells in the papilla territory (**Figure 5A**). This is just before the refinement of *Foxg* expression into the three cell clusters, or “spots”, corresponding to the future protrusions (**Figure 1**). At the initial tailbud stage, *NK4* appeared to be most strongly expressed in *Foxg-*negative cells surrounding the future papillae, including in *Foxg+* cells that nonetheless downregulate *Foxg* later on (Liu and Satou, 2019), as well as in cells of the presumptive ventral papilla (**Figure 5A**). *NK4* appears to be expressed at lower levels in more dorsal and lateral regions, especially near the two dorsolateral “spots” of *Foxg* expression corresponding to the two dorsal papillae.

**Figure 5.**
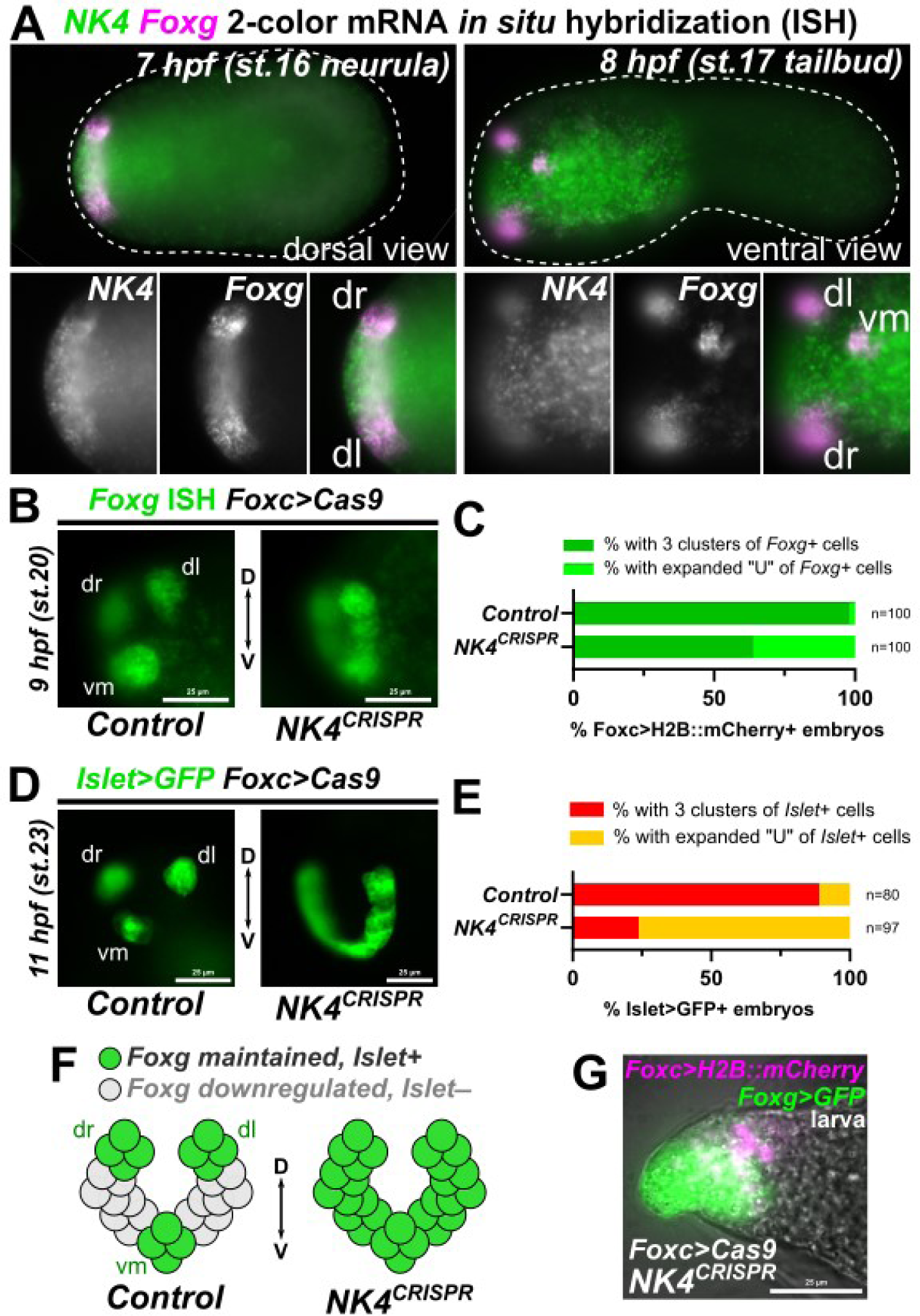
*NK4* CRISPR expands *Foxg* and *Islet* expression. A) Whole-mount, two-color fluorescent mRNA *in situ* hybridization showing *Foxg* (green) and *NK4* (magenta) expression in the papilla territory. Left: dorsal view at neurula stage, showing that *NK4* expression does not extend dorsally or laterally beyond the dorsal limits of *Foxg* expression. Right: ventral view at tailbud stage, showing strong *NK4* expression in ventral areas including the ventral *Foxg+* cells that will give rise to the ventral medial papilla. hpf = hours post-fertilization. This is a different focal plane of the same embryo imaged for a previous publication (Johnson et al. 2020). B) *In situ* hybridization of *Foxg* at 9 hpf (mid tailbud), showing expansion from three spots to a “U” shape swath of cells upon *NK4* CRISPR, compared to negative control CRISPR embryos. C) Scoring percentage of transfected embryos showing expanded *Foxg* expression as shown in panel B. D) Expansion of *Islet>GFP* reporter expression at 11 hpf (late tailbud) upon *NK4* CRISPR. Control embryos were instead electroporated with the *Defcab.47* sgRNA, targeting a gene that is not expressed in the papillae (see **materials and methods** for details). E) Scoring of transfected embryos showing expanded *Islet* reporter expression as seen in panel D. F) Summary diagram explaining our interpretation of expanded *Foxg* and *Islet* expression in *NK4* CRISPR embryos. G) *NK4* CRISPR larva showing *Foxg* reporter expression (green) in the resulting single large papilla. dr: dorsal right; dl: dorsal left; vm: ventral medial; D: dorsal; V: ventral. All scale bars = 25 µm. All GFP sequences are tagged with Unc-76.

Due to the fact that Foxg promotes Islet expression, and that both are required for the formation of the three protrusions later on (Liu et al., 2023; Liu and Satou, 2019; Wagner et al., 2014), we hypothesized that NK4 might downregulate *Foxg* in cells between and surrounding the three *Foxg+/Islet+* cell clusters. Indeed, we found that *NK4* CRISPR results in ectopic *Foxg* expression, observed by *in situ* at the mid-tailbud stage (**Figure 5B,C**). Instead of the three characteristic *Foxg+* cell clusters, we observe a “U”-shaped stripe of *Foxg* expression. This phenocopied the expansion of *Foxg* in BMP-inhibited larvae previously reported (Liu et al., 2023; Roure et al., 2023). We further observed that NK4 CRISPR similarly expands Islet reporter expression from three dots to a U-shaped stripe, like the expansion of Islet observed under BMP inhibition (**Figure 5D,E**)(Roure et al., 2023). It is thought that the expansion of these three *Foxg+/Islet+* clusters into a single contiguous group (**Figure 5F**) results in the enlarged protrusion seen in BMP-knockdown larvae (Liu et al., 2023; Roure et al., 2023). Indeed, *Foxg>GFP* reporter expression was observed throughout the single large papillae of *NK4* CRISPR larvae (**Figure 5G**). In contrast, NK4 overexpression (Foxc>NK4) suppressed Foxg reporter expression (**Figure 6A,B**). Taken together, these data suggest that NK4 is necessary and sufficient to repress *Foxg* in the papillae. Curiously, *Foxc>NK4* does not appear to suppress *Islet* reporter expression (**Figure S4**), suggesting more complex regulatory outcomes downstream of Foxg.

**Figure 6.**
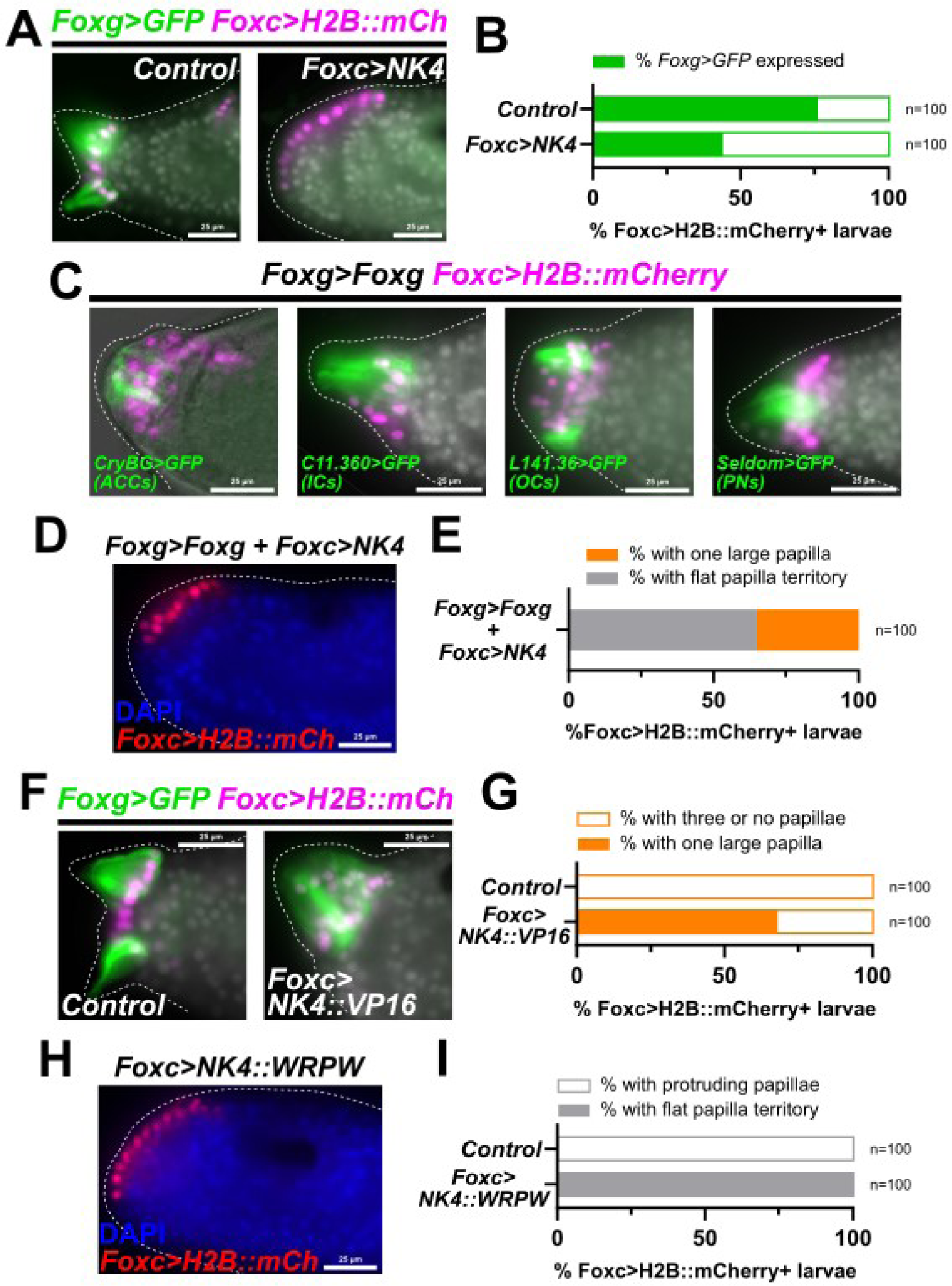
NK4 acts as a transcriptional repressor to suppress *Foxg* expression in the papillae. A) Overexpression of NK4 suppresses *Foxg* reporter expression (green) in the flattened papilla region. B) Scoring of larvae showing *Foxg* reporter expression, showing reduction upon NK4 overexpression compared to negative control larvae. C) Overexpression of *Foxg* by electroporating the *Foxg>Foxg* plasmid results in a single large papilla containing all four cell types, similar to that seen in *NK4* CRISPR and Noggin overexpression. OCs imaged at 21 hpf (St. 29) due to later onset of *L141.36* expression. ACC experiment conducted in a *Foxg* CRISPR background and imaged at 19 hpf (St. 28) as part of a rescue experiment described previously (Johnson et al., 2024), see **Supplemental Sequences** file for details. D) Single large papilla elicited by Foxg overexpression can be suppressed by co-overexpression of NK4. E) Scoring the percentage of transfected larvae showing either a single large papilla or a flat papilla territory upon overexpression of Foxg and NK4 combined. F) The C-terminus of NK4 (C-terminal to the homeodomain) was replaced with a VP16 transactivation domain. Overexpression of this NK4::VP16 construct resulted in a single large papilla with robust *Foxg>GFP* expression. G) Scoring the percentage of transfected larvae showing a single large papilla, comparing NK4::VP16-ovexpressing larvae to negative control larvae. NK4::VP16 also resulted in 30% larvae with no protruding papillae. H) Overexpression of NK4::WRPW, in which the C-terminus of NK4 is replaced with the WRPW Groucho co-repressor recruitment domain. This phenocopied full-length NK4 overexpression, suggesting NK4 normally acts as a transcriptional repressor in the papilla territory. I) Scoring the percentage of transfected larvae with either normal, protruding papillae or flat/no papillae, comparing NK4::WRPW overexpression to negative control larvae. All counterstained with DAPI (white or blue overlay) except *CryBG/*ACC image in panel C. All GFP sequences fused to Unc-76. All scale bars = 25 µm. mCh: mCherry.

NK4 has a repressive effect, whether directly or indirectly, on *Foxg* expression, and loss of NK4 impairs the refinement of *Foxg* expression into three discrete cell clusters. It was previously shown that Foxg overexpression results in a single large papilla protrusion (Liu and Satou, 2019). To test whether impaired refinement/downregulation of *Foxg* likely underlies the phenotype seen in BMP inhibition and *NK4* CRISPR conditions, we electroporated a *Foxg>Foxg* plasmid designed to overexpress Foxg and thus override the normal downregulation of *Foxg*. Indeed, the *Foxg>Foxg* plasmid results in a single enlarged papilla with all four major papilla cell types present, similar to both BMP inhibition and *NK4* CRISPR (**Figure 6C**). However, when we co-electroporated the *Foxc>Nk4* and *Foxg>Foxg* plasmids together in the same embryos, a majority of larvae developed a flat papilla territory, more closely resembling the *Foxc>NK4* phenotype (**Figure 6D,E**). This suggests that NK4 is able to suppress Foxg expression and/or function, and downstream papilla formation. Based on these results, we propose that the single, large protruding papilla seen with *NK4* CRISPR is due to impaired downregulation of Foxg in cells outside the three cell clusters that normally give rise to the three papillae.

NK4 and its orthologs in other species have been described as both transcriptional activators and repressors (Akazawa and Komuro, 2005; Choi et al., 1999). To probe whether NK4 is acting as an activator or a repressor in the papilla territory, we replaced the C-terminus of NK4 with either a VP16 transactivation domain (Campbell et al., 1984), or a WRPW repressor domain (Fisher et al., 1996), and expressed either throughout the papilla territory using the *Foxc* promoter. Strikingly, the effects of overexpressing NK4::VP16 appeared similar to the overexpression of Foxg, resulting in a single, large *Foxg+* papilla (**Figure 6F,G**). In contrast, overexpression of NK4::WRPW phenocopied that of the full-length NK4, resulting instead in a flat papilla territory (**Figure 6H,I**). These data suggest that NK4 normally functions as a transcriptional repressor of *Foxg* expression during papilla morphogenesis.

### Msx interferes with NK4-mediated repression of Foxg

While NK4 appears to mediate the downregulation of *Foxg*, we next asked how the ventral, medial protrusion forms. This protrusion arises from an area with high levels of BMP signaling (Roure et al., 2023) and NK4 expression. Indeed, by *in situ* hybridization, strong *NK4* and *Foxg* expression overlap in the ventral/medial papilla (**Figure 5A**), which is known to express *Msx* (**Figure 4F**). We therefore hypothesized that Msx might somehow interfere with NK4’s ability to repress *Foxg.* To test this, we co-electroporated equal amounts of *Foxc>Msx* and *Foxc>NK4*, to see if the former could neutralize the latter’s repressive effect on *Foxg* reporter expression and papilla formation. Indeed, co-electroporating *Foxc>Msx* and *Foxc>NK4* partially rescued *Foxg* reporter expression, resulting even in a large single protrusion in ~25% of cases (**Figure 7A-C**). This suggests that Msx can interfere with NK4 function, which may explain why the ventral papilla is able to form even in the presence of NK4. However, because the rescue was modest, there may be additional factors contributing to the de-repression of *Foxg* in the ventral papilla.

**Figure 7.**
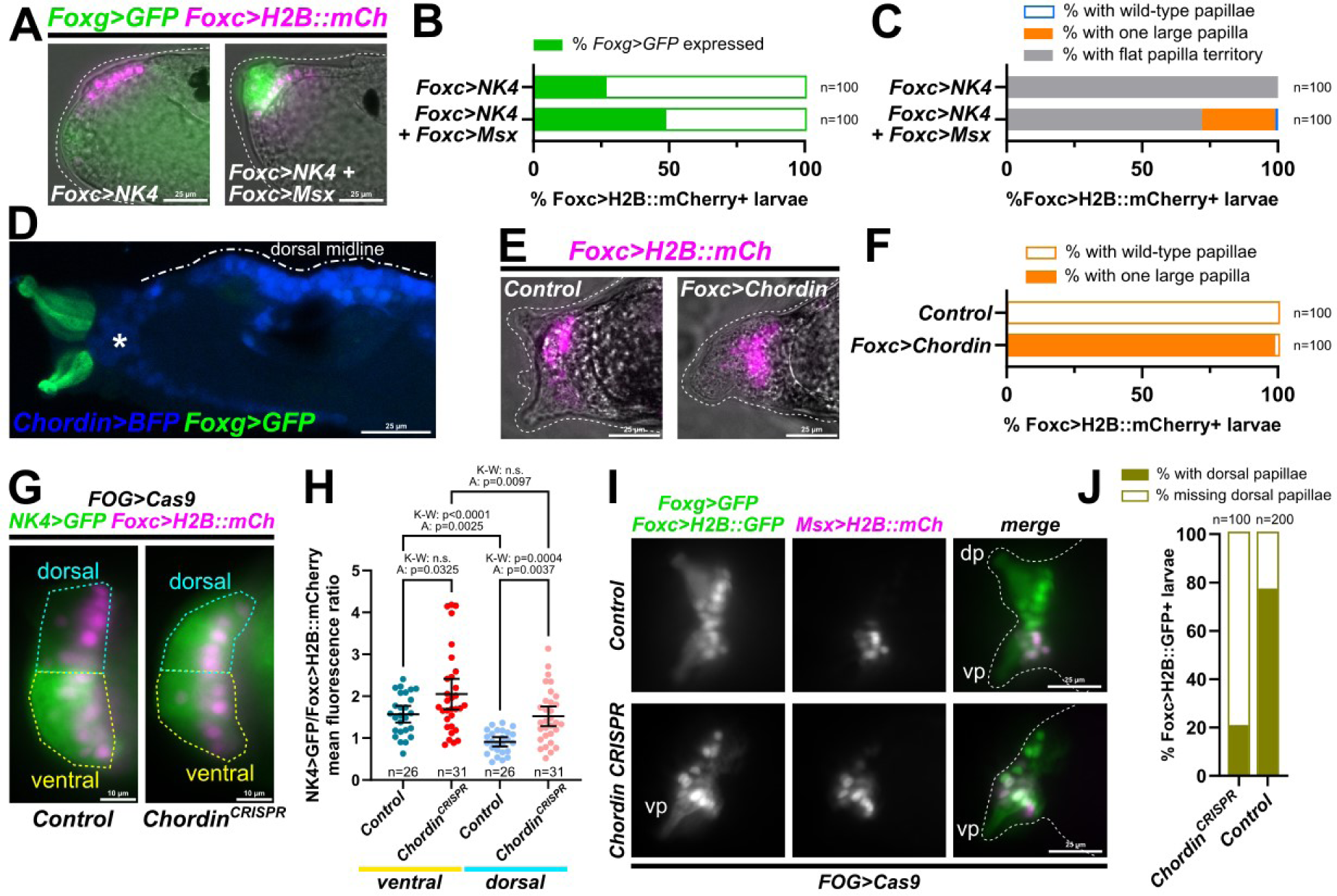
Regulation of NK4 expression and activity by Msx and Chordin. A) Overexpression of the ventral papilla-specific regulator Msx can partially rescue the NK4 overexpression-induced loss of *Foxg* reporter expression and papilla formation B) Scoring the percentage of transfected larvae showing *Foxg>GFP* reporter expression, comparing NK4 and Msx co-overexpression vs. NK4 overexpression alone, as shown in panel A. C) Scoring the percentage of larvae showing a flat papilla territory, a single large papilla, or the normal three papillae, comparing the conditions in panels A and B. D) a larva co-electroporated with *Chordin>TagBFP* (blue) and *Foxg>GFP* (green) reporter plasmids. *Chordin* is expressed in dorsal midline cells just posterior to the dorsal papillae (Liu et al., 2023). Asterisk denotes migratory mesenchyme cells, which also express *Chordin* at the tailbud stage (Imai et al., 2004). E) Overexpression of Chordin in the papilla territory phenocopies Noggin overexpression, suggesting it antagonizes BMP signaling as predicted. F) Scoring the percentage of transfected larvae showing either wild-type three normal papillae or a single large papilla, comparing Chordin overexpression and negative control larvae, as in panel E. G) Examples of the papilla territory divided into ventral and dorsal halves and imaged for *NK4>GFP* (green) and *Foxc>H2B::mCherry* (magenta) quantification, divided into dorsal and ventral halves, to compare *NK4>GFP* expression in negative control and *Chordin* CRISPR larvae. Scale bars = 10 µm. H) Ratios of GFP to mCherry mean fluorescence intensities were calculated for ventral and dorsal halves, across control and *Chordin* CRISPR conditions as shown in panel G. One-way ANOVA and Kruskal-Wallis tests were performed in parallel as the data for ventral papillae in *Chordin* CRISPR was not normally distributed on account of a few outliers. See text for discussion of possible interpretations. n.s.: not significant (p>0.05). I) *Chordin* CRISPR can eliminate the dorsal papillae, which are labeled by *Foxg>GFP* (green cytoplasm) but are negative for *Msx>H2B::mCherry* (magenta). dp: dorsal papilla(e), vp: ventral papilla. Scoring of larvae expressing *Foxc>H2B::GFP+* (green nuclei), showing presence or absence of dorsal papillae in control or *Chordin* CRISPR conditions as seen in panel I. All GFP sequences fused to Unc-76 except otherwise noted. All scale bars = 25 µm unless otherwise specified.

### Chordin restricts NK4 expression and allows for dorsal papilla formation

Having found that Msx can interfere with NK4 function, which may explain how the ventral Foxg+ protrusion forms, we next sought to understand how the two dorsal protrusions are allowed to form. By *in situ* hybridization at the mid-tailbud stage, we could not rule out *NK4* expression from the dorsal papillae even though expression was weak and did not extend more dorsally (**Figure 5A,B**). We hypothesized that *NK4* might be downregulated at the very dorsal edges of the papilla territory, enough to allow for sustained *Foxg* expression in the cells that eventually give rise to the two dorsal protrusions. Notably, the BMP antagonist Chordin is expressed in the dorsal midline of the epidermis, immediately posterior to the dorsal papillae (**Figure 7D, Figure S5**)(Imai et al., 2004; Liu et al., 2023). We hypothesized that Chordin might be able to limit *NK4* expression more dorsally through its ability to antagonize BMP signaling. Indeed, when we overexpressed Chordin (*Foxc>Chordin*), we obtained a similar effect as seen with Noggin overexpression, in which a single papilla protrusion formed (**Figure 7E,F**). In contrast, CRISPR/Cas9-mediated knockout of *Chordin* in the ectoderm significantly increased *NK4* reporter plasmid expression in the papilla territory (**Figure 7G,H, Figure S6**). By quantifying normalized mean *NK4>GFP* fluorescence, we demonstrated that its expression in negative control larvae is indeed lower in the dorsal half of the papilla territory than in the ventral (**Figure 7H, Figure S7**). CRISPR knockout of *Chordin* increased *NK4* reporter expression, especially in the dorsal half of the papilla territory, closest to the source of Chordin (**Figure 7H**). As expected, *Chordin* CRISPR suppressed dorsal *Foxg* reporter expression and dorsal papilla formation (**Figure 7I,J**). Based on these data, we propose that Chordin limits the reach of BMP-activated *NK4* expression on the dorsal side of the embryo, allowing for proper specification of the dorsal papillae.

## DISCUSSION

Here we present evidence for the involvement of NK4 as a key mediator of BMP signaling in patterning the sensory-adhesive papillae of tunicate larvae. It was previously proposed that BMP is required in the ventral papilla for the expression of an unknown, secreted factor that activates the expression of Sp6/7/8 (also known as Zfp220), which in turn was proposed to repress *Foxg* in interleaving cells to produce the tripartite organization of the papilla organ (Liu et al., 2023; Liu and Satou, 2019; Roure et al., 2023). This was partially based on lack of clear phosphoSMAD immunostaining outside the most ventral part of the papilla territory (Roure et al., 2023). Here we show that the missing factor may be NK4, and that *Sp6/7/8* need not be invoked. Instead of being secreted, NK4 is predicted to act cell-autonomously. We suggest that NK4 expression is activated by low levels of BMP signaling, beyond the limits of robust phosphoSMAD detection, which is consistent with our data and that of other studies tightly linking *NK4* expression to BMP signaling (Christiaen et al., 2010; Roure et al., 2023; Waki et al., 2015).

We propose that the patterning of the papillae can be explained by a classical morphogen gradient (BMP) with *NK4* and *Msx* as targets, possibly at distinct activation thresholds (**Figure 8**). We show that NK4 is a transcriptional repressor that is necessary and sufficient to repress *Foxg* in the papillae. A ventral-to-dorsal gradient of BMP signaling activates strong *NK4* expression throughout much of the papilla territory but not at the dorsal-most extreme, where we predict that BMP signaling is limited by dorsally-expressed Chordin. Here, *NK4* appears to be downregulated by Chordin, allowing for *Foxg* expression to remain active, resulting in the left and right dorsal papilla protrusions. In the ventral-most part of the papilla territory, where BMP signaling is at its highest, we propose that Msx interferes with the ability of NK4 to repress *Foxg*, contributing to the formation of the ventral papilla protrusion. Yet NK4 represses *Foxg* in all other cells of the territory, and loss of BMP or NK4 results in ectopic *Foxg* activation throughout and a single, large papilla (the “*Cyrano*” phenotype). Future work will be needed to confirm some of these predictions, and to ascertain which regulatory connections are direct or indirect, but this model is consistent with the data presented in the current study.

**Figure 8.**
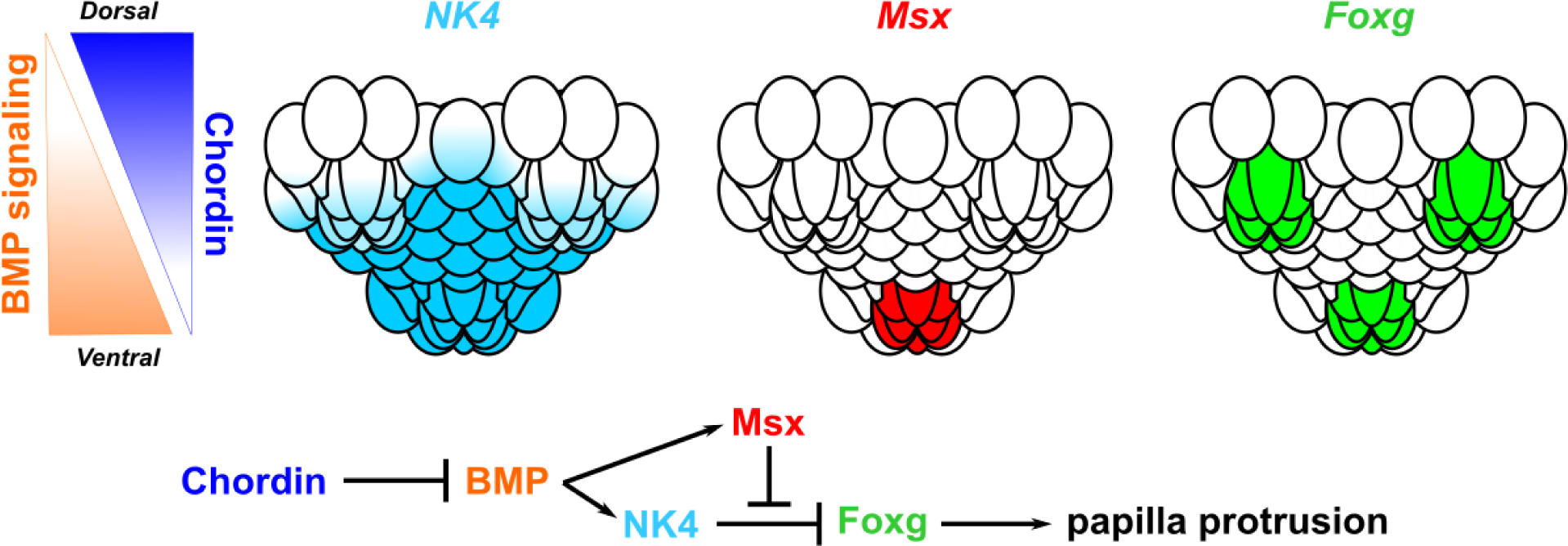
The proposed role of NK4 downstream of BMP in the papillae. A diagram summarizing the main points of the current work. Top left: A ventral-to-dorsal gradient of BMP signaling is shaped in part by an opposing, dorsal-to-ventral gradient of the BMP antagonist Chordin. Top right: expression domains of *NK4, Msx*, and *Foxg* in the papilla territory (dorsal at top). According to our proposed model, NK4 downregulates *Foxg* in certain intervening cells, but Msx may interfere with NK4 function in the ventral-most cells. Bottom: proposed regulatory network in which NK4 downregulates *Foxg* downstream of BMP signaling.

While our data strongly support a role for Chordin in antagonizing BMP and *NK4* expression in the dorsal papillae it is still unclear if this activity extends to more ventral areas. In our *Chordin* CRISPR larvae, different statistical tests were split on whether the increase in *NK4* reporter expression was statistically significant in the ventral half of the papilla territory, compared to that in negative control larvae (**Figure 7G,H**). Similarly, statistical tests were split on whether knocking out *Chordin* results in more uniform *NK4* reporter expression along the dorsal-ventral axis of the papillae. Accurate measurements may have been further hindered by the obvious morphological defects in the dorsal papillae elicited by *Chordin* CRISPR (**Figure 7I,J**). Although it is still not clear whether Chordin may affect the gradient of BMP signaling and *NK4* expression across the entire papilla territory, our data suggests that the dorsal areas are more sensitive to loss of Chordin function than the ventral areas. It is also unclear why *Foxg* is normally downregulated at the very dorsolateral edges of the papilla territory, where presumably NK4 should not be present to repress it. Here, regulation of *Foxg* may be independent of BMP and NK4, as suggested by the lower sensitivity of *Sp6/7/8* expression to BMP inhibition in these cells as previously reported (Liu et al., 2023). Similarly, NK4 might not be needed to repress *Foxg* in the very center of the papilla territory, which is derived from cells that never activate *Foxg* to begin with (**Figure 1**)(Liu and Satou, 2019).

One outstanding question is how to reconcile this model with data supporting alternative models. For instance, the MEK/ERK pathway (presumably downstream of FGF) was previously suggested to limit *Islet* expression to the three spots/protrusions, with pharmacological inhibition of MEK at the late neurula stage (stage 16) resulting in an expanded domain of *Islet* expression similar to that seen with BMP inhibition or *NK4* CRISPR (Wagner et al., 2014). MEK inhibition also suppresses the specification of Islet-negative cell types, such as outer collocytes and papilla neurons (Johnson et al., 2024). It is possible that MEK/ERK signaling might provide a temporal, not spatial, restriction on *Islet* expression. We know that MEK/ERK signaling is active in the lineage at a slightly earlier stage to promote *Foxg* expression in papilla precursors (Liu and Satou, 2019). If MEK/ERK signaling can repress *Islet* expression at this earlier stage (when its upstream activator, Foxg, is already present), it would explain why *Islet* is normally not expressed in a U-shaped territory like we see for *Foxg* early on. In other words, this might constitute a time-delay mechanism to ensure that *Islet* is only expressed in three dots, after *Foxg* expression has been refined. Future work would be needed to investigate this, with carefully timed *in situ* hybridizations to see if *Islet* is indeed precociously activated upon MEK inhibition. Another unresolved question is the exact role(s) of Sp6/7/8 in patterning the papillae. Previously we have shown that Sp6/7/8 is required for the proper specification of collocytes in the papillae, but this appears to complement, not repress, Foxg-mediated gene expression (Johnson et al., 2024). Furthermore, Sp6/7/8 is expressed in papilla neuron progenitors, but is not required for neurogenesis (Johnson et al., 2024). Neither BMP inhibition nor *NK4* CRISPR appear to prevent the formation of collocytes or papilla neurons, suggesting that more work will be needed to study the precise regulatory relationship between Sp6/7/8 and Foxg.

Finally, our model relies on some kind of interference with NK4 function by Msx. How might this work, molecularly speaking? Both Msx and NK4 belong to the NKL subclass of homeobox transcription factors (Holland et al., 2007). NK4 orthologs have been shown to dimerize (Kasahara et al., 2000; Zaffran and Frasch, 2005), and the residues in the homeodomain required for dimerization (Kasahara et al., 2001) appear to be shared between *Ciona* NK4 and Msx (**Figure S8**). One possibility is that co-expression of Msx and NK4 forces the formation of Msx:NK4 heterodimers that are unable to repress *Foxg.* Alternatively, Msx simply competes for NK4 binding sites in the *Foxg* promoter. Future work will elucidate the molecular basis for the ability of Msx to relieve NK4-mediated repression. In our experiments, Msx co-expression only partially alleviated the repressive effect of NK4, suggesting there may be additional factors expressed in the ventral medial papilla working together with Msx to promote *Foxg* activation there.

## MATERIALS AND METHODS

### Ciona handling and electroporation

Gravid adult specimens of *Ciona robusta* (*intestinalis* Type A) were collected and shipped by Marinus Scientific (Lakewood, CA) or M-REP (Carlsbad, CA). Gametes were isolated from adults and fertilized *in vitro* and dechorionated as previously described (Christiaen et al., 2009b). Electroporation of dechorionated, synchronized zygotes was carried out as previously described (Christiaen et al., 2009a). Plasmid DNA electroporation amounts were calculated and listed as total micrograms per 100 µl, for a final electroporation volume of 700 µl (including D-mannitol solution and sea water). DNA sequences and amounts are listed in the **Supplemental Sequences** file. Embryos, larvae, and juveniles were reared at 20°C, and staged according to hours post-fertilization (hpf) and Hotta Staging (Hotta et al., 2020). To monitor metamorphosis, larvae were transferred at 17 hpf (Hotta Stage 27) to a new agarose-coated 10 cm petri dish with 400 µl of 100X penicillin-streptomycin stock solution (Gibco) added to each dish. Juveniles were collected at 44 hpf (stage ~35) and fixed according to the same protocol as for larvae. Larvae were collected and fixed at 17 hpf, unless otherwise noted.

### Immunofluorescence, *in situ* hybridization, and imaging

Larvae and embryos were fixed as previously described for whole-mount mRNA *in situ* hybridizations (Stolfi and Levine, 2011) or for immunofluorescence or fluorescent protein imaging (Vedurupaka et al., 2025). *In situ* hybridizations were carried out as previously described (Stolfi and Levine, 2011). *Foxg* and *NK4* probes were previously published (Johnson et al., 2020). Acetylated tubulin antibody staining was performed modified from a published protocol (Pennati et al., 2015). Briefly, larvae were fixed by direct addition of 16% paraformaldehyde (250 µl for a 10 cm dish), then washed four times in PBS + 0.1% Tween-20. Larvae were incubated at 4°C overnight in mouse anti-acetylated primary antibody (Sigma-Aldrich catalog# MABT868) diluted 1:200 in PBS + 1% BSA, then washed four times in PBT. Secondary antibody (donkey anti-mouse AlexaFluor-488, ThemoFisher catalog# A21202) was incubated at 1:400 in PBS + 1% BSA again overnight at 4°C. Final washes were performed as before, prior to incubating with DAPI and mounting. Images were acquired on Leica DMIL LED or DMI8 inverted compound epifluorescence microscopes, or a Zeiss LSM 900 laser-scanning confocal microscope. Images acquired on Leica microscopes were processed and exported using LASX software, while confocal images were processed using the ZEN software suite. For scoring, only embryos/larvae expressing *Foxc>H2B::mCherry* in the papillae were scored, unless otherwise specifically noted.

### NK4 reporter fluorescence quantification and normalization

Fixed, slide-mounted larvae were imaged on a Leica DMI8 inverted compound epifluorescence microscope using the same illumination and exposure settings between larvae (though settings changed depending on whether it was GFP or mCherry fluorescence being imaged. For the Noggin overexpression experiment, the 20X objective (HC PL FLUOTAR L, 0.40 N.A., 0-2 mm correction collar) was used to image both the papillae and most of the larva (excluding the tail tip). For the Chordin CRISPR experiment, the 40X objective (HC PL APO, 0.95 N.A., 0.11-0.23 mm correction collar) was used to image mostly the head including the papillae. After image acquisition, regions of interest (ROIs) were drawn in Leica LASX software, and mean fluorescence pixel intensity values exported for further analysis and plotting. For the Noggin overexpression experiment, an ROI around the papilla region and another ROI around the whole larva was drawn for each individual. For the Chordin CRISPR experiment, one ROI was drawn around the ventral half of the papilla region, and another around the dorsal half of the papilla region, for each individual. Raw pixel intensity measurements and calculated GFP/mCherry ratios can be found in **Supplemental Data S1**.

### Tissue-specific CRISPR/Cas9-mediated mutagenesis in F0

CRISPR/Cas9-mediated mutagenesis in electroporated embryos was carried out as previously described (Popsuj et al., 2024). The original *NK4*-targeting sgRNA (*NK4.3*) was previously published and validated by in-house Illumina amplicon sequencing (Gandhi et al., 2017). A negative control (“Control”) sgRNA targeting no *Ciona* sequence was previously published (Stolfi et al., 2014). An additional sgRNA (*Defcab.*47) targeting the *Defcab* gene (Gibboney et al., 2020; Kim et al., 2025), which is not expressed in the papillae, was similarly used as a negative control in one experiment (**Figure 5D-E**). New sgRNAs targeting *NK4* and *Chordin* were designed using CRISPOR (Concordet and Haeussler, 2018; Haeussler et al., 2016), and validated by Amplicon E-Z sequencing service by Genewiz/Azenta (South Plainfield, NJ) as previously described (Johnson et al., 2023; Popsuj et al., 2024). The validation protocol can also be followed at dx.doi.org/10.17504/protocols.io.14egnr97yl5d/v1 (Popsuj et al., 2024). Cas9 was frequently fused to Geminin^N-terminus^, which increases mutagenesis efficacy (Pennati et al., 2024; Song et al., 2022). All sgRNA, Cas9, primer, and relevant plasmid DNA sequences and amounts are included in the **Supplemental Sequences** file.

## Supporting information

Supplemental Sequences

## ACKNOWLEDGMENTS

The authors would like to thank Dr. Florian Razy-Krajka for cloning parts of the *Chordin* promoter and coding sequences, and Dr. Robbie Richards for advice on statistical analyses. The authors also thank Dr. Eduardo Gigante and Hussan Ali for cloning and characterizing the *Noggin* coding sequence. The authors are grateful to other past and present members of the Stolfi and Cota labs for feedback and suggestions. This work was funded by NIH grants R35GM158421 to AS and R01HD104825 to CDC and AS, NSF (IOS) grant 2052493 to CDC, and an NSF Graduate Fellowship to CJJ. The authors declare no conflicts of interest.

---

**Supplemental Figure S1.**
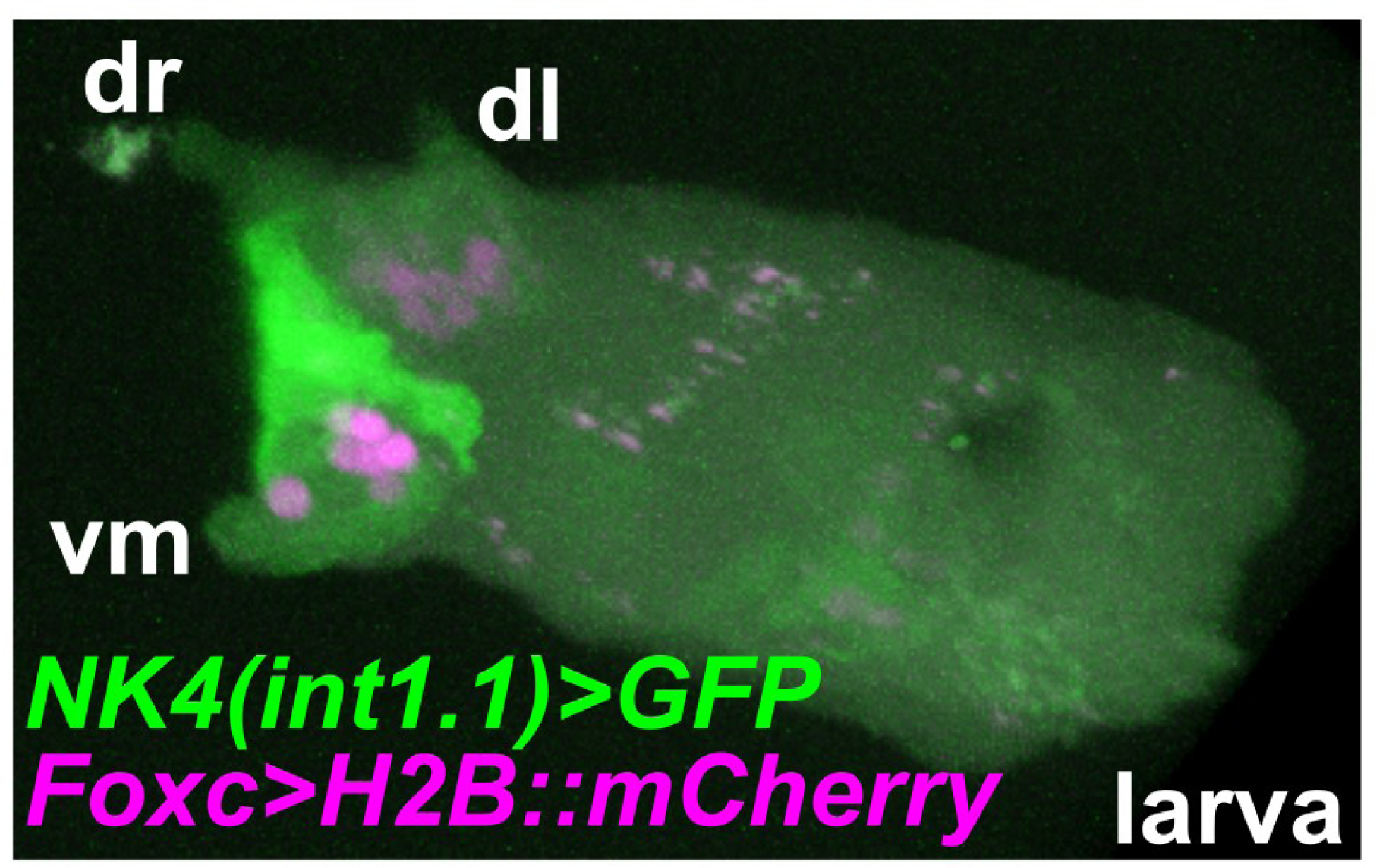
3D confocal projection of the head of a larva showing the *NK4* and *Foxc* reporters labeling the papilla territory. Note strong *NK4(int1.1)+bpFOG>Unc-76::GFP* expression (green) in cells located in between the three papillae, as well as in the ventral medial (vm) papilla, though it appears weak if not absent from the dorsal right (dr) and dorsal left (dl) papillae.

**Figure S2.**
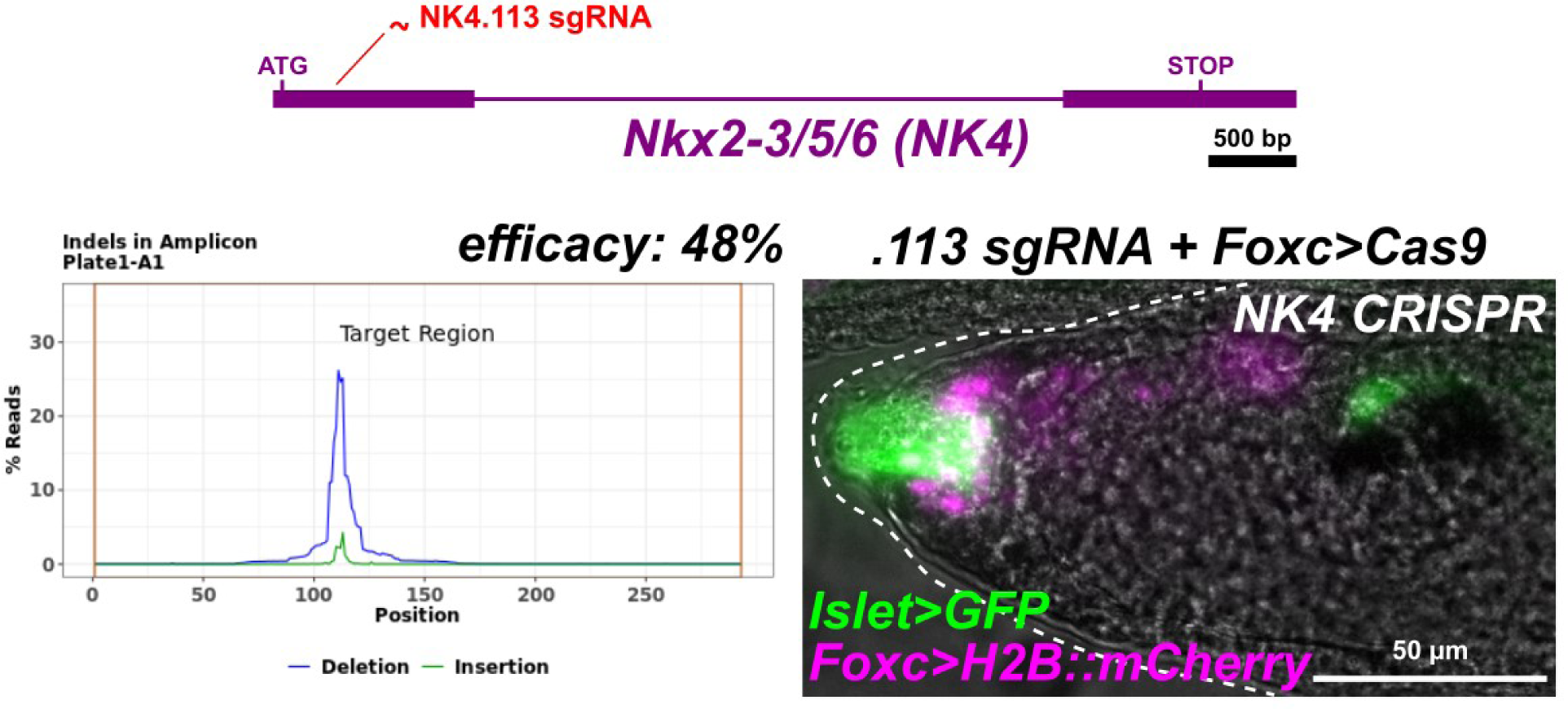
A second sgRNA (NK4.113) newly designed to target the first exon of *NK4* also results in a single large papilla. *Islet>GFP = Islet intron 1a + −473/+9>Unc-76::GFP.* The efficacy as measured by Illumina amplicon sequencing was 48%.

**Figure S3.**
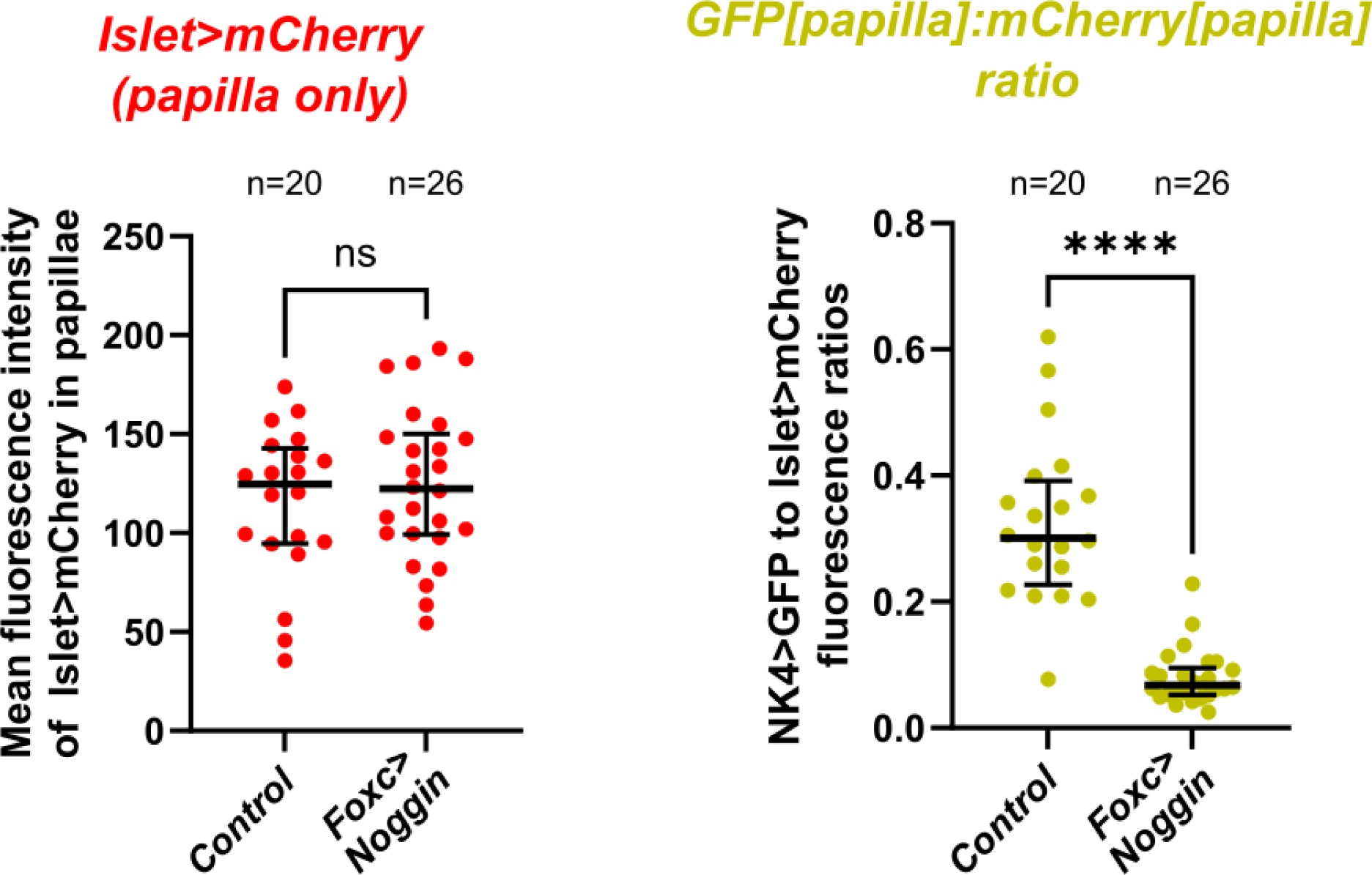
Comparison of mean fluorescence intensity readouts of *Islet>mCherry* fluorescence in the papilla territory only (unlike in main Figure 4D, which shows quantification of mean fluorescence intensity of *Islet>mCherry* in the whole larva), between negative control and *Foxc>Noggin* conditions as shown in Figure 4. Left: data show no significant effect on Noggin on Islet expression levels in the papilla, similar to how there was no significant effect on Islet expression throughout the larva. p = 0.6613 by two-tailed unpaired t-test. Right: ratio of *NK4>GFP* and *Islet>mCherry* mean fluorescence values in the papillae (unlike in the main figure 4E, which showed the ratio between GFP in the papillae and mCherry in the whole larva). These data show that the *NK4* reporter is significantly downregulated by Noggin overexpression even when normalized by papilla-specific mCherry levels. p<0.0001 by two-tailed Mann-Whitney. Outliers with GFP value > 80 (5 in control condition) or GFP:mCherry ratio >1.0 (two in the control condition, one in the Noggin condition) were removed from the entire dataset. Complete data with outliers flagged can be found in **Supplemental Data S1.**

**Supplemental Figure S4.**
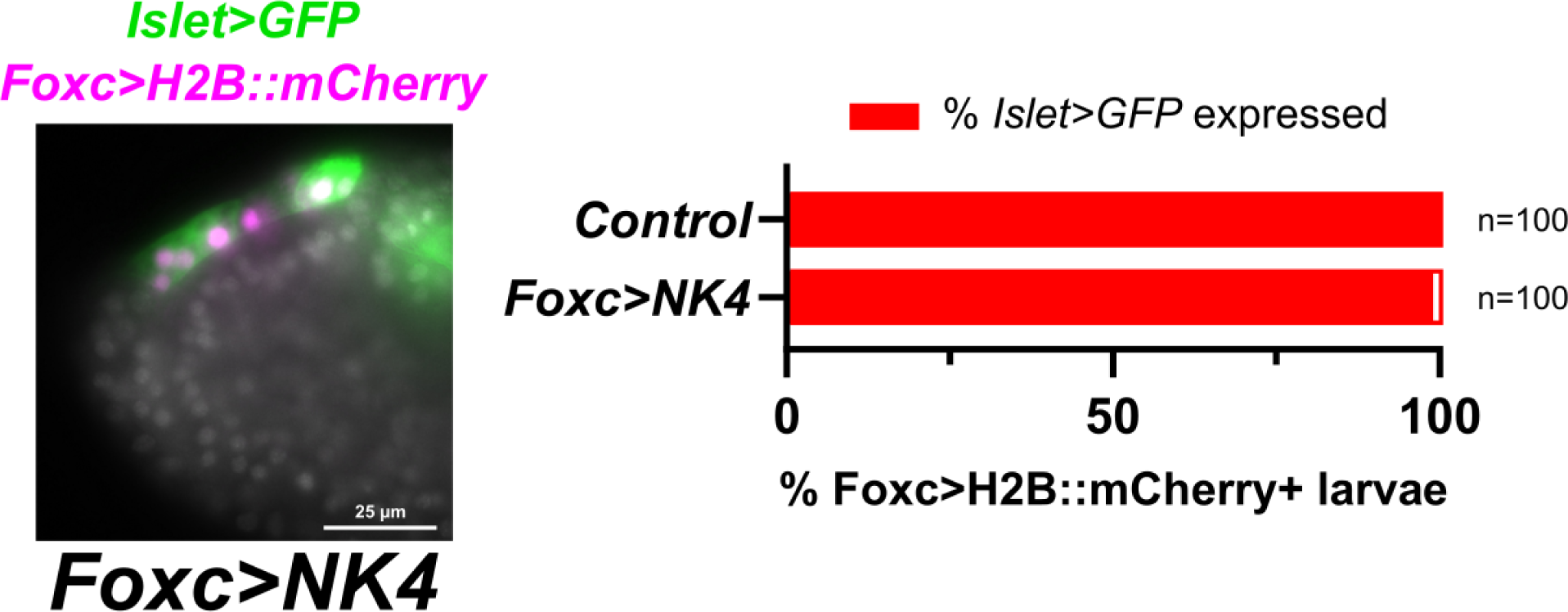
Overexpression of NK4 flattens the papillae but does not abolish *Islet* reporter expression (green) imaged at the larva stage. Larva is counterstained with DAPI (white overlay). *Islet>GFP = Islet intron 1a + bpFOG>Unc-76::GFP*.

**Supplemental Figure S5.**
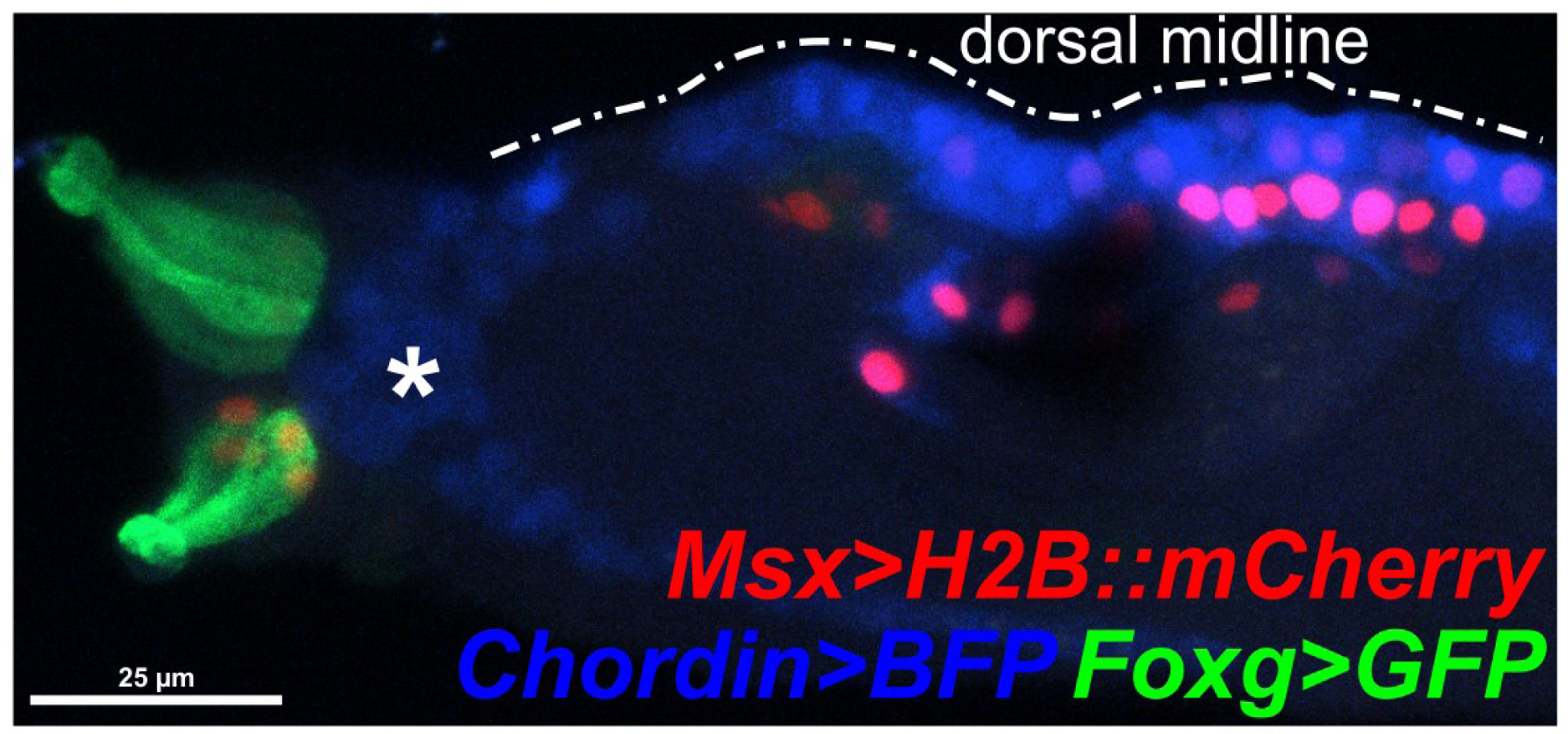
Triple-color image of the same larva imaged in main Figure 7D, here including also *Msx>H2B::mCherry* (red) which labels the ventral but not dorsal papillae. *Msx* reporter expression can also be seen in the dorsal midline. Asterisk denotes Chordin+ migratory mesenchyme cells. BFP = TagBFP; GFP = Unc-76::GFP.

**Supplemental Figure S6.**
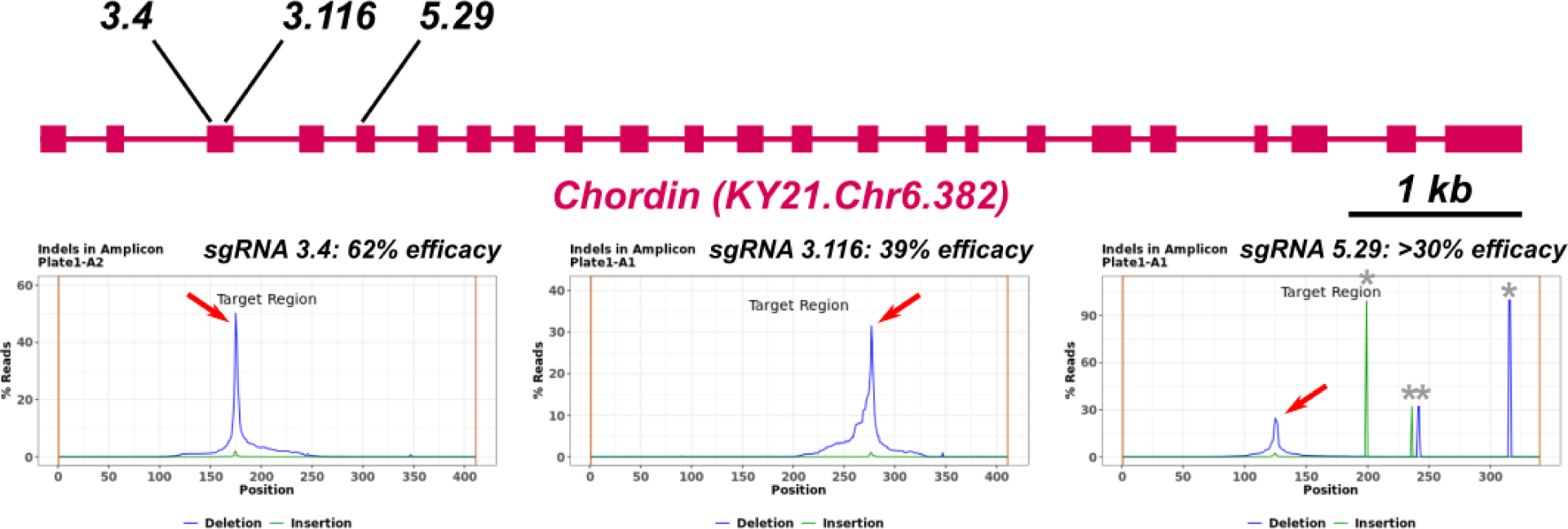
Validation of three sgRNAs targeting *Chordin*, following the protocol in Popsuj et al. 2024 and at: dx.doi.org/10.17504/protocols.io.14egnr97yl5d/v1 Red arrows indicate CRISPR-generated indel peaks. The validation estimate for sgRNA 5.29 is not precise due to the presence of naturally-occurring indels (grey asterisks) in the amplicon sequence.

**Supplemental Figure S7.**
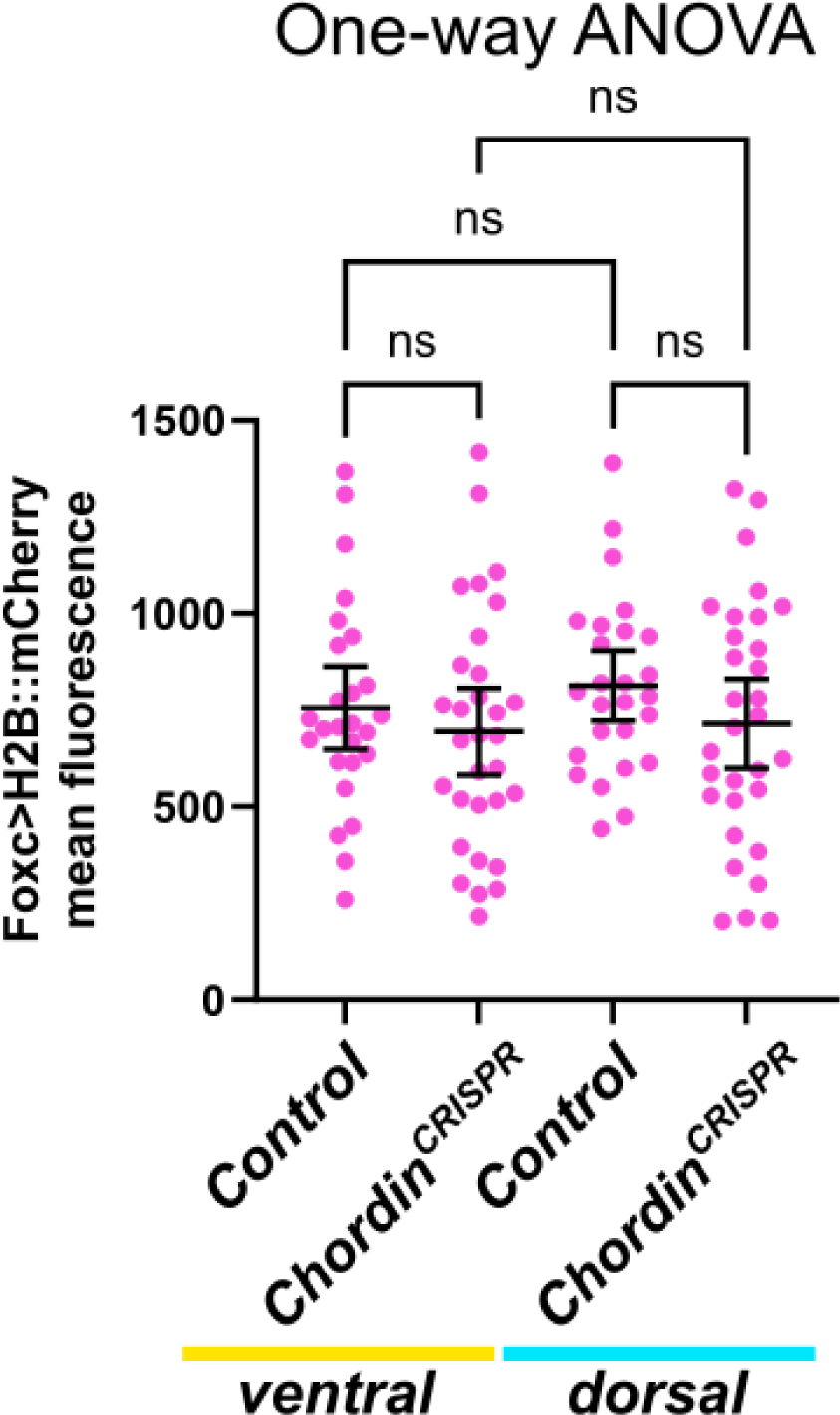
Quantification of *Foxc>H2B::mCherry* mean fluorescence in negative control vs. *Chordin* CRISPR larvae, subdivided by ventral or dorsal papilla territories. These are the mean fluorescence values for mCherry underlying the ratios plotted in main Figure 7H. The mean fluorescence values are not significantly different between the different halves and/or conditions, suggesting the differences observed with the ratios of GFP to mCherry mean fluorescence were indeed attributed to changes in *NK4>GFP* expression. See underlying raw data in **Supplemental Data S1.**

**Supplemental Figure S8.**
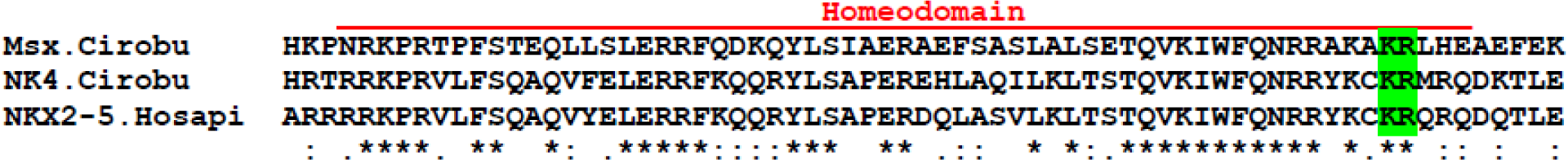
CLUSTAL O(1.2.4) multiple sequence alignment of the homeodomain of *Ciona robusta* (“Cirobu”) Msx, NK4, and Homo sapiens (“Hosapi”) NKX2-5, showing the conserved KR motif mediating homodimerization of NKX2-5. See **Supplemental Sequences** file for full protein sequences.

## Supplemental Video descriptions

**Video S1.**

3D projection of confocal image Z-stack through a larva co-electroporated with *NK4 int1.1 + bpFOG>GFP* (green) and *Foxg>H2B::mCherry* (magenta). Ventral medial papilla on the left, dorsal to the upper right.

**Video S2.**

A Z-stack series from confocal imaging of an *NK4* CRISPR larva (using *Foxc>Cas9::Geminin^N-terminus^)*, co-electroporated with *Seldom>Unc-76::GFP* (green cytoplasm) and *Foxc>H2B::mCherry* (red nuclei). Apical cilia are labeled by immunostaining for acetylated tubulin (AlexaFluor-488, also green), showing that PNs surrounding the single large papilla are still ciliated. Larva is counterstained with DAPI (blue). Anterior to upper left, dorsal on top.

**Video S3.**

3D projection of same larva as in Video S2.

**Video S4.**

3D projection of a confocal image stack through a negative control larva electroporated with Seldom>Unc-76::GFP (green cytoplasm) and Foxc>H2B::mCherry (red nuclei). Apical cilia are labeled by immunostaining for acetylated tubulin (AlexaFluor-488, also green). Larva is counterstained with DAPI (blue). Anterior to bottom right, ventral on left.

## Notes

### Competing Interest Statement

The authors have declared no competing interest.

### Summary of Updates

Now includes Illumina-based amplicon sequencing-generated indel plots for validation of NK4.113 and Chordin.5.29 sgRNAs.

