## Supplemental Sequences for "An NKX2-5 homolog is required downstream of BMP signaling to pattern the sensory-adhesive organ of a tunicate larva"

**Supplemental sequence file – NK4 paper**

Sequences might be based on genome assemblies, and might not be verified by plasmid sequencing.

>NK4 intron1.1 + bpFOG

(basal promoter of FOG from Rothbächer et al. 2007)

cacgggtaagttgttttacttgtctcgttgctttaaaagcagcttacgcatgaataagcggaatgggaattagctgaataagatctacgagcctttatgaccctgactatatttagtaaataaaactcctggaccagcgcgttatattttgaaggcaggtagacgtgtcaagtttcactagcgtgagtcggtttaacaaaatatacttaatgcgaagcggagcgaatcgcgggaccaggaaacaaatgccgtgaaccgtgcagggtacaattaacaagcgatacttgacatgtggtttggcatagtttagcggcatattagcaccaaaggggcacccaaacaaatcacacgagcgactttccatgcagttatgggcgacagtgacagacaaagccacaactcggtggcgccggcgttacgacttagaagcaggtcggccaatcacaacaaaaagccacagcgatggagatcggtgctaatgaaaacatagatgcgagtaatgcgttacaataacacggggcgaggaatcaggcagtcggcgactatcggaaacggctccagtgaatcgaagccccattgtcccagagcgcgacaagcgcctcgacaaacaagagaacaagagataattacaaattaacaagggtggcgatgtatggcgggccgtgtcgccgaattcaaatagcgcccacttgaaatgatcgcggtaattttggatagactctgtcgccgccattttgattatcttgttaagatttcaagtgtcggttgcatggtaagcttgttgaccaaaacaatatcgttagcggattacttgcacattggtcttgtttcgataaacaaacaactaaatcgatgcaaaatttagtgtcagtgcagcgcacacaaaaaaacgaatccgtctttccgcgaagctgtaaatgttgaaattcctttccataaaccgtaactggtcatattacgagaagtacatagagagttcaactcgctcaaaaactgtaaacacttgcgagtaaccagcaattataaccaaccgcaaaacgaagtgtctcgagtatctgcaggtcgactctagaggatccggcaaagcttcgtgtattgtaccggcccattgtcaatcatgcaaacttgatattatattgacaagagaagaaggcagtttaaattaaaactctaaagtagagagacattaatctcagctgacaaggcaggtggtcacagtaagttcatttaaatagttggccaacaatagcctttccaagaaagtatttttgttccaggtctatacaaaaataacacacaacatg

>Foxc -2132/-1 (Based on Wagner and Levine 2012)

ccgcctacgtagggtaaataccgctcagggcgcgtacgttgcacacagcgggaaatgacaaagaaatggataaaaagatgcatggttttctaaattgtccggcatacagaaaacattcccctgcaaagttatatacgtggtaactcgtaagtgggaatgcggtgttataaaacaaaacacccatgttataacgaccgtcgttttctcggcacttgataataaataaatcgcattcattcaattaataatagatagccagcagcacgcgtatgttatttgagcaatcgcttgcaagaggcacgtccgggaagtccacaccttctacctgaaacgtctactttgacgttatgaacccgtgcaccgaaaagtgttagcgaaaccaagatggttaaactggaatagcttcctcaacacccacgtacactctttccttttgcgtcaagcggggtttcctttccctctagaagtgttgatataaatccttcatgacaccggacaccgcattaagcgcggttcaatcagaatatctatgcggcacccgcttcatgtagtaggccgggggaagatgggacacctttagcacataatacccaaacaacctaatcgtattttaaacagttaacaacggtctatggaagtcgtgaggatacggtttaataattctttaaatatttcttgtttactaccaaatgagacgagaaaatagaataaaaaagtgtcccatctccccccaccctactgtataaaccagaaaaagtggaaatgtccgaaaagagttttatcagcattttagtattttggcgatttaagctttagatacacaaggtgttagtaattgcggaaaggtgttttgttcagttaatctgacgaaaggagttggtttatatttttatttctaccaatatatatatattgctaccaggatttagtaaaaagcgttttatagttaatttaaaagtaaatattttaaacagtaagcaacaaatctgtacgattaattgctccgtaaacgtttactgcatgttcggttaaataaccacctcgcttttcatttttcaatcctactcgttaacgtcgttttgctaattccctttacttttttcaaggccaagttagttactataccaaacgcaatataacataacttaaactgttgtttagctatttaaatacagaaacaaaaataagattaattgaaatagcaaaccaagagaatcagcaacaaaactacacttgttaaaaatacgctatgaaggtaaaaaaaaactaagtaaaaatgtccaatatatttaataaaacgaagctatggtgggtggggaaaccttaagctaaatgctccaggaaaatatgaatcatcgacgcctaagttgccgccttagttgcataactcattgtatagcgagtcacggaaaatgctgcgcgtgtaaatttccgcatggtgtcgctgctgagccaaccggtctcgctcgtttcaaaaagtcgagttttaccgcaaaaaactcttcggttacatctcttatttataaacagcaacccgaggagtcacgctgtaaactgatcgggtcgtgacaaggttcggacaggagaggcagcttcagttataaccgctgaatatcaacggtgaacgttaaccgccatttttaatgaacgttggagttaaaaagttccaagattgagagattaatttaaaagttgtggtttatataaacagggctattgggtaaggctccatagtgagcggtgtcagcaggtgtttcgtaaggcggcgcgtgccaagttctctacttagagcttgtcaaaacacgatctaattactgcatcattagcgcgccattgttcctcgcgaaagttgattgggattatgacgctcctgctttccattgtttaaggggaagatgaactttttaccttcgctcaggctcgactcggtcgtgggcaggtaccggcagaaaacattcgattattgacacgaaggcagtgcgagtgttgtgagggaagtcgtttcggagcgacgtttgtttgcttgcagcgttggcgttcagattctaacttttatatatctcgggcagtgttagtgtaagttaagttacgttgaacacaggacaccgaatccttggtttgattctctata

>CryBG -1068/-24 (Based on Shimeld et al. 2005)

taattcttactgttcggttgaaactcttgaatccactatgacgtcatcatcgtcgctacaaccgctataaacagcgagaaacaaaataaaaacaatcgacttcattacggaagagttgggtcgaaagtggttgttgcatgaaatatatgtgtgttatatctcggtgctaagtggtacaacaaacaaataaaacgtctggtttatgagatgctcgcgtggtacgggctatgtgatgtcataatacctttgtctattatccttgttacgacataattaccatgaaacagcattgtgacgtcacagtgctgtttgtgttggacaaacaacacgtggtttctacgagggggaatcccctctctgtcattattacgtatgatgacgtataatatcgattattattcaccatccaaggttatatacgatatctatttggttttatccgcttggtggcgacatcaatcctcaactacaattggtattatgacgtagttcgtagttacgtcacaagtcacacaacaaagaatttattatgagcaaataaacgaactaatatgacgcaacaataactaacaatttatcacgtgatcgttttatttattttgtttgttatgtcataaccacttggtgcttaaaagggagcttatgacgtattaatgaactttaattattaaaacaatcttgcttatcgtgctgtgttacgtcataatacatgcattagcaattactttgttttgtcataatatgaattatgtaattatcgtgacgttgtttgttttaaaaacccatttattattacgtcataatactttagatggttgtgtatgttacgtcataatgttaattgttgaataaaaatgaagaaacaaagccatgaagaaggcggtttgttataaaacattgttgaagggatggaattaattatgttgttaacgtttgaattattgatgtaacaaggacgacaagtgagcgcaaatcagatttcgtttactttccttctaaccctcctaaccactgcttatttcgcattttgtacaatcgaagtttc

>Seldom -2991/-12 (From Johnson et al. 2024, originally KH.C4.78)

ccatgatcgctgcaagtttaggttcgccgtttcgcactcatgttaccgaaccacctaacctcattacactggtgccatgaatctgtaaagttgtaatttatctgtcaccaaaaatgcccttacgttataggtaaacgggttatttagctataaaagaagtttttgcttgcggccgtcgtaaggaatccttgaccgacgtataataaagtttcttttcattttttaatatgtcatcgtaaaaacaaggatgaaactgtcgtaccacagctaaactcctatttttgccaatacaggtaaaatctcctgaaagtaggactgttttgccaattgatttaggagtcggacctacgcactaaaccgcgcttatatgtgccagcataatacactgcctttcaaagtcgtatcaacagaagcctatccctaaaggtggaattagcaaagcaaatattatctctgacgcggcctggtagcccttcttttacagacggtcaacattgtctttcaaataataataagtaagcgcaaatacgaatcagatggatgacttagtgctgccatggatataaaatattgttctgcataagttgataaatataaagttttggtttctaaaacagaagtaagacttttttaatttttatattacaaagtgttaataatgttcgccgttcagtgtaaatatgggggcggggggaaacgggacaccttcagcagataatatccgaatatcataaataacacagagtaagatcacactttacgtcaactaccaaaagttaaaggacaattagtccctagttatttagtaggttggggggagataggacacgtttaacacattttattcaaatatcttgatcgtgatataaacaactaacaacgttctatgagagtagcggggatacggttttataattatttgaatgtttcttgtttaccaccaaatggaacgagaaaatagaatgaaaacgtgtcccatcttcaccctgctgtcggttttaactttataataaaaactatttttccgaatgaccgaccgtcgcgcatctttactacgacgtcggttaaaaacttttttgctacatagttaacataactatcttcatgtatttcgagataatgagagataatatatcgccatagcttgttcatgaagtaacatgtagacaaacgagactactcaactcccgtaacctcacaaactgcaaacacttatgtcgtttaagtatgtaatcaattgtactgtgactttcggttaatgtaacgcttcgcctatagacctacagcggttaagtaatatctaatttcgggttaaatttgccgaatccaatatattcgtaaacgtgactcccagactgtcgcaagaagtcaatataaatatctcagtggccctgtaataattctaaatcgagttgaatcaatttggttcgcaaccttatcgcttcttattctggataatctaatcactggcatttgtgtacgctgtttgtgggcgaggaactcgctggcctttctataaaaacacaactcacaaagacctggcaagcggcgtcttttgttgctaggtcacattcagaccaccgcgcgcggtatacgtgttgcttttttctagtacagccaaagcgattctcttcaaaatacagtcgagctacacttcaattaaattcactttcaagcgaatctacatcgaaactaatgaattttcaacattaattagtagccaagtttgaaactaataggtcacacttgtattttggttagtttttatccgaacctttttgaagacttttttggctttcaacgataaataatatgccgagagacgttttataatgaatccaataaagaactcaggcttatattagactaccaaccctaacagttcatgaatcatttatataaggataacccgaatgtatacacacctatacgtatttaaaaacagctcaccactttttcagcgtaacatgtttaggcctagtttattacagctttgactaaatttgacttcacctttgattacatgtcttcaaattttggtcgttttcacagtaatagaatttcgataactccaaacgaatcagcaacataaatatgcgggcctcactcgtctggggcacaactcgtatgcaaagtttaaaactacgttcaaatgttaacttgcatccggattgtgacgtcagagaaaaacgtaagttacgttttttgcttgtctatatttgttaaagcgtcacgtttttattcaatttgatgaaaacaatgagcgattaggtttagaaaagcaaacacgtatagataaattttgtttttttgaccaagtcaacgaaagttaaaatagtatatacataaaaaattttgacacgctaaaaaggtataaaacttggatacagaacgtaggataaccgttatacaatgagaaattttattaatgtaacattattgggaaactgtggaaaaagagtttagacacatgactagtacaataatgggaactaccaaatttattttttttactaatgaaaaaatattatcacgaatgaattacgtaaacgagttttctccatacaattaatttcagatcaaaagcggaacggatagtcgtcattgcattaataattcatacacttggttgacttatacccgtcgtcaacatcaataattgatcttaactgtcgcggacataataaattcggttgctcgaattttttaaggtaattgactcccgtgttaaggtatgcgaatgttggctgtttagtgtctgctagcgtggaaataaattctttctcatgtttgctggttttcagtatagtaaattagggagattggaatttgttaaataagctgaatgtgtaaatttagtcagtaaatttaaataggatgtaatgtgtctaatacctaatacgttgtcgattattcagcgactttttaatgatgtaaatttcagacatttattaacacggacaaaaacttagtactga

>KH.C11.360 -1395/-3 (From Johnson et al. 2024)

gacgtcatcatggtacaaagacagtcaggcgttttatctgtgacaaggatcacacgtcatcagctcgactggttatacaattaaaatcgattcgcagcgtaatgacctacataaacccgcagtgataacacattagctatgaatggattcacgctgtgtttgaatacataaccctagttcaacgtacagacacggcgacggcgtacgattataattcaaactcgtaaaacgttccaacgcgtgtgcgctattccataaaatttctaacttataatcgcattataattgagcacatttcagtgcgccgttgtaactggtcgctaataacaagcgttgttcttatttaaggtcttggtatgtgttcgctactgtgaactctgctcgactgatacttctgactcttcacaacctgttatatacacgcaccacagtcagtgttaaatgtaacacatttcatccctagcgcctcgcgctcgtccaacttcttttcgctttttcatgtcggcgtttgaaagcgtcggcgctcgagttagcaaataaggtctttcttttcacactcggcaccagtccagggtaggagagatactacagaccaataatccgcacgagtatcatcaccttcgctaaacaagttctctaattcggggcatgagacgctggctacggttttagagcgtcttttgtttcaggtgcgccttgcttcgaaggttattgagataacgctgtgcactaatccacaaaacctttaaatctactaaacaaatgtaaatagtttagaatgtctcttttgcgttttggcttgcaacattattttattgttatcaaaacatatggactacaatcacaacaaacaccgtaagccgccgcttattaatccatttttgatgatcagttttcgttctaattcacgaactcaattaatataaatgtttaaaccaaataagtctacattttgcgttgttgtgtaaaatttattctcttcgcagtttcaattaatataatcgtcgtaaatcatatatgcgacacccaaaagaaaatttgtgtcaaccacaatatgaaaaaggacgcatataagaaccttatacacattgcaaatccattatggcctgttgtttttaaagttttaattgctacattatacgatacaaaacatacgcccaaatgcgtattgctacgtacatgccgacctattgccaatatatttagccaactaaatgcatatcgtaaagatttaataacatatcatacggatttgacgtatagtttcttttatatgttacgtaacccttagtacataagacttcactttcaaagactgctatgttgtgtggtttaattctagctccgtcatactgaaattttattttctaaactaaacgaagtttttcttgt

>KH.L141.36 -3401/-1 (from Johnson et al. 2024)

aaccactgcggttttcattatattaccaaacgcttatttcatacgatttgatcttaaatggaaaggaatgtttgaaaaatagagatttgctctgacgtcattaatttttttgacattttgtcagatttgcttgtgttggtgtattgatcttataatgtatcatggtggagggttagtactaaggataggaaatattaagtttataaataaatctgtaaacctttgaggtgtttctatttatactatatcgcgataaatatttacattttttcctttgtacagtggcccatgtatttaataccttgcaacaattcaagcgagacaaccaaggggcttatgcacgcattgcgtggtacacagttattgcgattacttgctgcaaaaatacaaggtcacagcatcaccacacaaaccactacaaaaaggaatatcttatgcaaatttaaattgttgtgtggctaattgttcaaaacacgattagaaaacagataatatggcatttaggggctaacggtatcccatcttaccccacagactgctgtgtaataatctgtacaaatatgacagaatagcgatttattattttaccaatatacatatttcatattattacaaagcactatagtcatacttagtaccgatgttttattccttgattctaatactcactgtaggggaaatccaggtaatttgttactgccacgaaacttaacacttacatactgacaacttggttcacacgcgtatctgcctatcgaagatctgtttcattattacacggaccatttgttcagcgacaccaaaaccctttgactttcctgagattacaggtttgaaacattgccgttaaatgatctccaccaaccgggtacaaggctcatgcttcgccgccatatagcacgcttcaactattacaacgcccggtgacgttggtcacgtgcctgtctcgcacttggcgcttcatagaggctaggcttcagtgagatactcgattgcgacccggggtatatggggtgcattgcgcgacttcgtgcgtacattcttcctcgttggcggcgaatgaaacttttgtcgtggccatatatgtttaaaatcaacactacagcaataactggtatacgataaaacaatataaatctttatttgttttgctttcaaacaaacgaagatcagattaattaaaacgaagggagggaaattaacagcaatacgaccattgtttaaaacaaggtacgaatatgttgtatacttcaatgaaagtgtcaaataaaacatgtcacagcatatattgtagcaaaaactgtgacgatcaaatatacagagaggaatgacttagataaagttcagttatatctgtaatatcaatgctaatcacataacaaagttgtacgttgtccaccattggtatactcacttacagacaatagttatctcccgaggtttttcgtgcaatttgactgcatttctttttattgttaatcaagtacgaattctgggcagaataacttgtagcaaatacggcatgttgtaatttcaagaaagccgttaaaagacagcctgccccagttctatttagaacgatacattttcgagaacctccttaaccgttaaatatttgtgtattaggctattgctaaaattattttcttacagttttgaaacgaataggttttttatcgtataaaaaatttaatgttaaattatttttcatatttttactgtttaaagttttagcgattattccatgagtcccaacaaacagcgatgatgcgactcgggacgcccacttgcggattgaagaaaccattctcttaaaaatagatatttgcaatgccccacggccgggtctgtcttttgtttctgataccagagaacccatgtcgatatattgctgatgacgtaacatggaatactgttagcacctatattccatattttctacttgtgttttggacaattgaaaacgttttatttacaacgctgttttagagcagtgggaataaggttatataattctgtaaatattcattgtttaacaagtgggacggtaaaagtaaatgaaaacatgtcccattttaccccaaccaatgacatcatttattatcaacactataattatatttactacatatagcgctgtataaagttgatgtatatatatctatatacatcaatataaattaaaacgtaattaagcaaagtataaaaatgttgtttcgtttttgcttttatcaattctacctcctgtgtttattactcagaaattcatgggagaaaattgacgcctctggtttttcccatttcgcacaaactacttctcagaaattatctcaatgagcatgtgactaaaaatctatgttcttataatggtatattttagttgaagtaccccttgggtttagatcactaaattacccgaaaaatttttttttaagtttgtccccaaaaaaactgcaaaattacaatgcggggcaaaatggcgaacggcatagaaaggttttgatatttaaaaccaattaaccctctttattttaattgctgcagataattattatatttaacattctgtcgattgaaaatatgcgggcgtgtacttaatggctataaattttaataaaaagagatatactgagatttggctctttcaattaggtgtttcacgcagactaatatttagttccgttcaattacaacaatattcacgttttatcgcaatcctaaatatgacattacgagacgtaatttaaattaagataatctgatcaaggagttcgaatagccggaggggggaatttactttaacgtaagcaagcctatatatattgttttgccaaactttcaaatttataattagtttagcggatataatgtgaaattggtatgtattaggttggggaaagatgggatcttcatgtttgggtagaacgggctatacgtgcccgctggggtaagacgggaaacttaaactggggtagaatgagacagctgggttatggtgagatgcgacgctgtatagaatgttcgattttcattcccttataatattttgtttgcttgtaaactaagaatatttacagaattatatgaacgtgtatcccaagactctaaaagtctccatctaaccctacaagactttagcgccgcgaatgtaaagacagtcgttaattgtttcagacacgaccaggaattatgggaacttgttgctcacaatatatcttatcccacagtatataggtataagttttaaaggaaccgtc

>Islet intron 1a (0 to 2014) + -473/+9 (mutated ATG) XhoI

(originally from Wagner et al. 2014)

gcctcgcttaattgcggtaagtttgtgggttgtttaataaagtaggggggtttgggttcgaaatcaggccagggttgaaaatccagggatcccgggttctaatccaggtcagggttggagaaccagggatcctgggttcgaattcaggccaagaactagcattgtacgcgtaagtgtcttgaacaaaattgcgggtaagtgcttaagcaggaaaggcaacaattaaccttaggattcagtatatagtgcgaagacggatacattattaatgggggggcgacctttgtaagcgcctaacattggactgggccggggggttggggcgcgatgttcgcgatgaaattggatttttatcttttggcaacaaaattaccgttcgcgtagatcgccatctttaaaccgcgcggcgaatatctcaagccattgtcacgtcacaatcggcatttattatttatttacgtcacaacggcttaatctgaagctgcaaaaggtatttgtgacatcacaaaacataacaatgaagttgtgttaaagcattgtgaatcacaaatacgagataatatacatcgcttatgtcgtcataaccgcgtcgatgcagccgcgtaaatcgcactcaatgtcccattcaagctgctacgtcacaatcgctattctttccccacaatagacaatgttagttaacaatttcccgtgtgtgacgcaacaattaaaacaataaagaacaatgcagtgtattctaatacattgtagaatgctcgcgtcatcttacagcgcacaccgcggttgcatcgaatgaataatgcaacaacatatcgacccgcattcgataaaatatcagacttttatattttccttaaagacctcaagtattttaactttttttcgaaaatcttttttattttaataatattttcgcagtgacgtcattaatttatgacatcaccagtgatatcatgacgtcactcgatcaatattagccaagagcaaaaatagtggatatttttcattaattctttatttgtttatttaaccggaaacaaaatggccgccggtgacctggttgattgtgacgtatataacacgcaacacacgactattatgacgaaacacagacttgaattgaaaatctagaaccccggtcacgttatggtaatgcttatgacgaaataattgaaagtttttgcgattgtgacgtcataatacctttattgttaggcgattggttaattaggacgcgtgttcgtttttaattgtgacgtaataaacatgaaattaaacgagattgccaaagttaaatgatgcattatgacatcataatatataacttttatgtgattctagaaaagacgaaaccgcgaaagtagaacaagacgattgttaacaaagctacgtcataatccataatcaacacgcgaatcgcgtcgctaagtatccgcacctgtgacgtaacaatactaagacacaggcaaacttcgatatcgctaaaatatatgatttctatgcgatcgcgattaagtggttttaattgttcgctagcaaagtggtttgtggagattaatttgaggtaatgtagtttagaaccaacttattgtaaattatctaaagtggtcacatgtatacgcacgtgtatatggattaattaaggggggtcattaagtgtaaagtcgctaatccccctcacccgtgagactttagtcattaacaaagtgtttgttaattgatcgacaaagtgggtcgcggtgtaatagttgctgcattatgacataacatgcattataaatgtgacgtcacattgaataaacaatcgtctatatttggattttaggacttgagataaaaaagttaaaatatgtttgggtcttgatggcttgtaacttttgttggggattattttgttaattgttgggagggtcaggtttttaagagctttgttaagcgttaaataaagaaagaaagaaatttaaataaattaatttaattttagattgaactttcttaaacacagaaataaaataaagtttttgtttggcctcgagggttaacttaacatgggcgtgtgaaatgtaattagttaatgtagttagtcgtatctggatcggaaggcattcatttgcactgtaggcattgaactaggaggttcaccacatcgaggaatatttggtcgcgctgcggggattgagggccttgtagttggggggaggtttgaaagtttcccgaagtaatattaactttgttgtataggacccgttgttgctgttagagagaatcggagaaaaagtttcgagagaagctttagataaattgattctgtcgcattcgagtattacaacttgtgtctgtgacgtgagcgagtgaggacaatttactgtgacgtcataatagccagcaatataggggtggaaggtgaataccccgctctacaagcggagaatttcgaaatttcaaacctggatcccggatactcgccgccgttggcggacgactcaaggtcggggtcgttcacaagttttatcaacgaa

>Islet intron 1a (0 to 2014) + bpFOG XhoI

(originally from Johnson et al. 2024)

gcctcgcttaattgcggtaagtttgtgggttgtttaataaagtaggggggtttgggttcgaaatcaggccagggttgaaaatccagggatcccgggttctaatccaggtcagggttggagaaccagggatcctgggttcgaattcaggccaagaactagcattgtacgcgtaagtgtcttgaacaaaattgcgggtaagtgcttaagcaggaaaggcaacaattaaccttaggattcagtatatagtgcgaagacggatacattattaatgggggggcgacctttgtaagcgcctaacattggactgggccggggggttggggcgcgatgttcgcgatgaaattggatttttatcttttggcaacaaaattaccgttcgcgtagatcgccatctttaaaccgcgcggcgaatatctcaagccattgtcacgtcacaatcggcatttattatttatttacgtcacaacggcttaatctgaagctgcaaaaggtatttgtgacatcacaaaacataacaatgaagttgtgttaaagcattgtgaatcacaaatacgagataatatacatcgcttatgtcgtcataaccgcgtcgatgcagccgcgtaaatcgcactcaatgtcccattcaagctgctacgtcacaatcgctattctttccccacaatagacaatgttagttaacaatttcccgtgtgtgacgcaacaattaaaacaataaagaacaatgcagtgtattctaatacattgtagaatgctcgcgtcatcttacagcgcacaccgcggttgcatcgaatgaataatgcaacaacatatcgacccgcattcgataaaatatcagacttttatattttccttaaagacctcaagtattttaactttttttcgaaaatcttttttattttaataatattttcgcagtgacgtcattaatttatgacatcaccagtgatatcatgacgtcactcgatcaatattagccaagagcaaaaatagtggatatttttcattaattctttatttgtttatttaaccggaaacaaaatggccgccggtgacctggttgattgtgacgtatataacacgcaacacacgactattatgacgaaacacagacttgaattgaaaatctagaaccccggtcacgttatggtaatgcttatgacgaaataattgaaagtttttgcgattgtgacgtcataatacctttattgttaggcgattggttaattaggacgcgtgttcgtttttaattgtgacgtaataaacatgaaattaaacgagattgccaaagttaaatgatgcattatgacatcataatatataacttttatgtgattctagaaaagacgaaaccgcgaaagtagaacaagacgattgttaacaaagctacgtcataatccataatcaacacgcgaatcgcgtcgctaagtatccgcacctgtgacgtaacaatactaagacacaggcaaacttcgatatcgctaaaatatatgatttctatgcgatcgcgattaagtggttttaattgttcgctagcaaagtggtttgtggagattaatttgaggtaatgtagtttagaaccaacttattgtaaattatctaaagtggtcacatgtatacgcacgtgtatatggattaattaaggggggtcattaagtgtaaagtcgctaatccccctcacccgtgagactttagtcattaacaaagtgtttgttaattgatcgacaaagtgggtcgcggtgtaatagttgctgcattatgacataacatgcattataaatgtgacgtcacattgaataaacaatcgtctatatttggattttaggacttgagataaaaaagttaaaatatgtttgggtcttgatggcttgtaacttttgttggggattattttgttaattgttgggagggtcaggtttttaagagctttgttaagcgttaaataaagaaagaaagaaatttaaataaattaatttaattttagattgaactttcttaaacacagaaataaaataaagtttttgtttggcctcgagcagctgaagcttgcatgcctgcaggtcgactctagaggatccggcaaagcttcgtgtattgtaccggcccattgtcaatcatgcaaacttgatattatattgacaagagaagaaggcagtttaaattaaaactctaaagtagagagacattaatctcagctgacaaggcaggtggtcacagtaagttcatttaaatagttggccaacaatagcctttccaagaaagtatttttgttccaggtctatacaaaaataacacacatagc

>Foxg -2863/+54 ATG start codon

(originally based on Cao et al. 2019, updated in Johnson et al. 2024)

gtttcattttccaacaaaacattataatgcgataagagaattggccatctttaccacagaacttaaatttaacggaaatataaagttgcaacgtacgtaaataacagaccacccgtatttattgcgtgacattaatacattggatgtcaaagaacaaagaacttgttgtttacgtcatataggatccggtacttgataaaccacttcaagaataagatttatgacatcataaccaaacatgagcgacgactttaattgggaaaaaatactttgaaatgttttttttatcagatatttgacaacaaaaatacaatagtcggaaaatattgtattttttagttttatttgttcatatttaatatataagcacacctaacacctggctgataatgggacgtcactgggttaaaatagaaatcatataaagcacaagttcaattggtttcgggctaaaaattaaatcctttttcggtaaaccgaccgtttgtcacatcaaagccacagcgagcagtgataagagccgagtgtttttatgtaaataggtttaggtaactcgtgtctgcttgatttggttgtaagattaattactcgcagtgttacacatgttcccacaaacccagagacattgtaactggggcgtagatgatacgttataaaatataaaccaacacatgttgtataatataagtgcactcattttatccgcagagtgttataacgattttcgcttttaaaatatatagtagggtgggggaagatgggacacctttagcacataatatccaaatgttctcatcgcgttttgaaccatttcaacggtctattgtcgtaaggatacggttttgtaaatctttgaatgttttttgtttactaccaagtgggacgagaaattagaatgaaaaggtgtcccatcttttcccgctctaatatataactttttaaatctgtgtaatgtttcatttaaagatgttctgggctattgagttcttaaaacttatcaaacgtatgttggtatatctgcccggacctttattattaaacaccaaaaccacatagaaaaagtaactattaaacgaaattctgtctatgtcagccgttaaactaacttgcagtgtgtatagatatcgttgggctctatttggtggtaaacaaagaatatttacagaattatataaccgtattcttacgactgtaaaagagtgttgttaattgtttaaaacacgaataaaaaatatgggagatcgtgtgctgatggttttcaccttatcacagttcagataaaaaattaaaagttctcaaaacaaactataggaacaaatatgtttttgatttatattattgcaaaaatactatgttaaattggcctacctatttccaattaaaaaaagtcacattttaattaattttacggaaacactgtacgttgtcagtgattttagacctgatcttataattgagtctggagtaaaacagaccttgtgattgacagtttgaattataagtatcagaaaacattgatcgtagatgtaaccaaagctctcatcagttgtggaccaaagtaaccacaaaacctacctaatttttgcagtgttgcctttagactaaacttttatttcagtgttgccatttaaaaaaaaagctagatgtgaggtcggtgaataaaaagggtaaattattagttatatgagcgtagtttaagctataactagttaatacagatatattaaatcatgcagtttgcgtacaaattatattaattaaagtccattaaaaaatgtattcaatattacaaaaaaatcaatttctattaagaaaaacagaaaaacattaaaaaattggaattttttgacattttaatcgtggcaacatgacaaacttatgttagcagattatcacgccatcttaacatacagcacgtgtgaagcaagatcaaataaccaaggtgtcaagtgaaagttacagggcacaattatttaaccaagctaacgtaaatgaaaaacccaggcaaccggcttatcactgctggctacttggtcagtgtggttaacatatatcaacggaaagccccgctctatatcatgacgtattaactagtattaacaagcccaacattgtacggtgtcgtgtaaaaccttatatggcatgaagcttgttttgcggaaagctattttgttttttaatctgaaaatatgagatgggtgtggtatgggaaaatgaacgcctatagacacacctggtgtattcacacaccttatcctcactatgttgagttatagcatgggtaattcttatccaccacacatggtgtatttagtgtattcatacttagttataacagtattgtatatttaaatattgccaaaatccgtcgtagagtgacaaaaacaatcttgaaatactgcaaagtcattaggcgtattatctcacactctaaatcacttaaacacaataaagtcgcgagttttaaaagcctagatctaataataattgcaaatcaaacgcattatgtccattagttgtattttctttgtcgatttagtatccgggttatccttgttaacaaggtgtttgtgtgtctaagtatgtccatgattacgtcttcctaccagaggagtcgtgttgttttagtcaatacggtggtaattatccgattattgtgatatagcgagttacaggcgccctcagctcacggtggaagtgctaacttgtaagtcttaatgcatttctcttttcttcaaaggtttttgagtagattttcgtttgtaattataccggcgtctccttacgctgtaatccacgcttttaattgtgacgtggattgacagaaagagtagcgcgagagagaaacgcggcatacagaatATGACGAACGACGCGGCCGAGTCTGGTCATTCAAAGCGAGAAAACTTCATAGAG

>Msx -2298/-73 (from Stolfi et al. 2015, based on Russo et al. 2004)

ctacgcattgatgtcgcaatctcaccaagtttcccaccacctatatgaaagctacgaaggcggaaagatttcggcataataacctattgttgacgtcataatcgaccgataaataagcggcgaagatcgcgtcctctacaataggaggtggaacactcataaaaaatcacagcggtaaagccgctaccacacgctgctaacatttattaaggagacatgctacactcgacgtcataatgcgcgtgacgactatattgtgacgtcacagagagataaataggatatttattgcgcggtagaatagaacgacgagcaataaaatgaaatgaattttggaaatggtaaattaatttattttataagtcgtctaaactttatccatttccgtatagaacacgacttttaccaactgttaaattttattaagttaccacataaagtagtttattgagccgctctgaacgaatggaaatcaccgtactctattaagttgatcaaccttttacatttcaatatgttaaaaatcgtacattaaaaaaagatatagtaggatggggggatgggatgcgttttcattctattttcttgtccaatttggtagtaaaaaaggaactttcaaagatttataaaaccgcatcctcacgactttcatagacggatggtaattgtttaaaacacgaacagtatattaagatattttctaatggcgtcccgtcttcccccgccctactatatattataaagaggtaatatgaaccaatatttagaattttagttggtgcacatcaattactgtttttataagtccatgcagaccgatgtccatttaccagtgattaaacagggcaataaaattacgaagaatcgccttgcgtcgctgcaaacatatttattaggcattagcaagtgtgacgtaataatgcctcttgttcgacgtgacggcgagcgcttgcttttgaaacaccgcgacgtcgacgaagacggtgtcgtaacttaacaatagtctggggcctatgatgtcataatgcccatttggtgcactcgcgtgctgtcgcccaaaatcgcaattaactcgatgacactggcagcatatgacgtcataaaatggttgccagggtcccggaccgagacagtaattcacttttatgacccgaatatctaatgggacccatgatgtacacttagaagcacgcagcaactgattattaatgttgagtgtgttacggatgactgttaatttcacaaacaaacaaaaaaacagagttcgttggcctttagatcactattagaaacaacactaattggtgggaacttgtttcggagcgagccaaataaaaaagctaaaacaaacataacagtaaagctttattaaaataataagcgttgttaataaatattagcatatagaccaaacaccaaattgtgtgcaatcagtttttgcatctttaaaactattaaatttttatgttcgccgtgcgaatttttatttgaatgattctattaataaagcctgaaatcctaacaatcattttgagcttccacatttgatattaaaattctcgtatttgttgttatttattaatgattttgagtgacttatttagactcagttttttttttttcaattattcaagttcggatgtgtggagatacgggattaatacaatgaatgttcaattaaaaacacaaaagttttaatttgaattaatgtcggcgctgcagacgagccgagagcttgtcggatttaaaatgtaatcctccaactttgtctggcactgacggtgtagatgcaatctgaaaatggcgactgagcaggatcgggtcgcggctggagctccggcgggctcgggacgctcataattccgccggtaatccccgtaacgtcgatgaaagcgaacgcgccgacaaaagtgacgaagattaagtgtaacaacatggtttaaacaaacagactggagcagcggagacgagagagagagggaacgtaatcgccgcggccgaagcgtgaccaagtcgttgtattgttataatagaaaatcgttttattatgatttaatgaacgcgcgctggcgactgaaggatgctgtataattgggggtaatggggaaatatcgggagaaaaaatatggcgatctgcgggagaggatagtcgcttacataatctctggagtttagttgaagcagttcatattgagagaaacatccctcagacgagtagttcgtaaa

>Chordin -1732/-43 (based on Abitua et al. 2015)

cggttacgttaagttcgtgcgtagttttacataacgtaactctgccaggattttgttacgtatatgttactaccaattaaagtgcaattagggatttatagccctgtggttcgtgcaaacttagtatacacacatgcatgcgtatagttgttcaatgcgtcaaaatagtgtcgatatttttttcgcggcaatgtcatattttaatgccatttatctttcaaagacggtctgccattttgggattttttgtccacatacggttaaaatatgtttattaatgttcaatttttttccttgtgatacagtttcataacctgtacgttttcttttaaaaaaaaggatctttttctgtcgtttttttcaccatttttttactcttacttgcaaaaaaacggcgtctttgcgtattttgcgttaaattatgtccaagtatttattattacacatttctaacaaggttatagacgttttgtggcgtgaaacgtagtagacacgtatagtgtttttgtgtatacgcctcttactgtcgatgcttactgttaagttataacattggcgttatacactgggacacatgatgtattcatacaccttatgcttactgtcaagttataacatgggcgtcttattcaccagacacacatggcgtattcaaaacagagctacaaggtaaacagtctgcatgatatactttaaccgactggagccttttctgcaaacattttcccaaactcggattcccacggttggcccgctattccattgtccttgcatacagacatctgttacgaaacggtctttgttgtgctattatgtcgaacacaatagcgtgtatgtctgttggtataattaggtaacggagcacgcgaatgctttattatgtaattagtaagacacgctcattaaaaattaagataaccgaccgtggcaattggtttaacttaaaaaaaagattattttaagaattcaattttacggagccaccatagacttaaataattagactttaaatgttgtgcctaaaatatattttacgcggtttttccgtggagcaagatggatactattaagcacataacattttataatcatgtttttaacaatcaacaacattttatccaatcttaacaatcaacaacacacaatcatgtttttaacaatcaacaacattttaggaatcgcgaggatacggttatatggatctgtaaatgttcattgtttactattaatagggcgataaagcgaatgaaaacatggaccatcttacctcagcctactataccacaatgacgtaataccttattcatttgtagatttactatctcacgtcaccaaacacatgactcggccaaataaattcacacttttataaaataaatattccaccggattctgcacttggctagctgcaagttcggaaacgagttctgttacgtcacagcatgtaatatgacgtcacagcaggtaatgtgacgtcgtattggtgcagacacgcgcggcctgcccagtttttgcccacaaaggccattgtcccgaacctgctatcccgcttcgaggctccagtcggaattgtgacgtaataatgaaccgtgacgtcacgggaaaaaggctacgtcacagtgggggtactttatctagggatctcgtagatttcaccatatcgaaatctcacttcgagcaggaacaaat

>Cas9::CionaGeminin-Nterminus (from Song et al. 2022, which originally described it as having the Geminin sequence from human instead)

nls::Cas9::nls (described in Stolfi et al. 2014)

Ciona robusta Geminin N-terminus

atggctagccccaaaaagaagaggaaagtggacaagaagtattctatcggactggacatcgggactaatagcgtcgggtgggccgtgatcactgacgagtacaaggtgccctctaagaagttcaaggtgctcgggaacaccgaccggcattccatcaagaaaaatctgatcggagctctcctctttgattcaggggagaccgctgaagcaacccgcctcaagcggactgctagacggcggtacaccaggaggaagaaccggatttgttaccttcaagagatattctccaacgaaatggcaaaggtcgacgacagcttcttccataggctggaagaatcattcctcgtggaagaggataagaagcatgaacggcatcccatcttcggtaatatcgtcgacgaggtggcctatcacgagaaatacccaaccatctaccatcttcgcaaaaagctggtggactcaaccgacaaggcagacctccggcttatctacctggccctggcccacatgatcaagttcagaggccacttcctgatcgagggcgacctcaatcctgacaatagcgatgtggataaactgttcatccagctggtgcagacttacaaccagctctttgaagagaaccccatcaatgcaagcggagtcgatgccaaggccattctgtcagcccggctgtcaaagagccgcagacttgagaatcttatcgctcagctgccgggtgaaaagaaaaatggactgttcgggaacctgattgctctttcacttgggctgactcccaatttcaagtctaatttcgacctggcagaggatgccaagctgcaactgtccaaggacacctatgatgacgatctcgacaacctcctggcccagatcggtgaccaatacgccgaccttttccttgctgctaagaatctttctgacgccatcctgctgtctgacattctccgcgtgaacactgaaatcaccaaggcccctctttcagcttcaatgattaagcggtatgatgagcaccaccaggacctgaccctgcttaaggcactcgtccggcagcagcttccggagaagtacaaggaaatcttctttgaccagtcaaagaatggatacgccggctacatcgacggaggtgcctcccaagaggaattttataagtttatcaaacctatccttgagaagatggacggcaccgaagagctcctcgtgaaactgaatcgggaggatctgctgcggaagcagcgcactttcgacaatgggagcattccccaccagatccatcttggggagcttcacgccatccttcggcgccaagaggacttctacccctttcttaaggacaacagggagaagattgagaaaattctcactttccgcatcccctactacgtgggacccctcgccagaggaaatagccggtttgcttggatgaccagaaagtcagaagaaactatcactccctggaacttcgaagaggtggtggacaagggagccagcgctcagtcattcatcgaacggatgactaacttcgataagaacctccccaatgagaaggtcctgccgaaacattccctgctctacgagtactttaccgtgtacaacgagctgaccaaggtgaaatatgtcaccgaagggatgaggaagcccgcattcctgtcaggcgaacaaaagaaggcaattgtggaccttctgttcaagaccaatagaaaggtgaccgtgaagcagctgaaggaggactatttcaagaaaattgaatgcttcgactctgtggagattagcggggtcgaagatcggttcaacgcaagcctgggtacctaccatgatctgcttaagatcatcaaggacaaggattttctggacaatgaggagaacgaggacatccttgaggacattgtcctgactctcactctgttcgaggaccgggaaatgatcgaggagaggcttaagacctacgcccatctgttcgacgataaagtgatgaagcaacttaaacggagaagatataccggatggggacgccttagccgcaaactcatcaacggaatccgggacaaacagagcggaaagaccattcttgatttccttaagagcgacggattcgctaatcgcaacttcatgcaacttatccatgatgattccctgacctttaaggaggacatccagaaggcccaagtgtctggacaaggtgactcactgcacgagcatatcgcaaatctggctggttcacccgctattaagaagggtattctccagaccgtgaaagtcgtggacgagctggtcaaggtgatgggtcgccataaaccagagaacattgtcatcgagatggccagggaaaaccagactacccagaagggacagaagaacagcagggagcggatgaaaagaattgaggaagggattaaggagctcgggtcacagatccttaaagagcacccggtggaaaacacccagcttcagaatgagaagctctatctgtactaccttcaaaatggacgcgatatgtatgtggaccaagagcttgatatcaacaggctctcagactacgacgtggaccacatcgtccctcagagcttcctcaaagacgactcaattgacaataaggtgctgactcgctcagacaagaaccggggaaagtcagataacgtgccctcagaggaagtcgtgaaaaagatgaagaactattggcgccagcttctgaacgcaaagctgatcactcagcggaagttcgacaatctcactaaggctgagaggggcggactgagcgaactggacaaagcaggattcattaaacggcaacttgtggagactcggcagattactaaacatgtcgcccaaatccttgactcacgcatgaataccaagtacgacgaaaacgacaaacttatccgcgaggtgaaggtgattaccctgaagtccaagctggtcagcgatttcagaaaggactttcaattctacaaagtgcgggagatcaataactatcatcatgctcatgacgcatatctgaatgccgtggtgggaaccgccctgatcaagaagtacccaaagctggaaagcgagttcgtgtacggagactacaaggtctacgacgtgcgcaagatgattgccaaatctgagcaggagatcggaaaggccaccgcaaagtacttcttctacagcaacatcatgaatttcttcaagaccgaaatcacccttgcaaacggtgagatccggaagaggccgctcatcgagactaatggggagactggcgaaatcgtgtgggacaagggcagagatttcgctaccgtgcgcaaagtgctttctatgcctcaagtgaacatcgtgaagaaaaccgaggtgcaaaccggaggcttttctaaggaatcaatcctccccaagcgcaactccgacaagctcattgcaaggaagaaggattgggaccctaagaagtacggcggattcgattcaccaactgtggcttattctgtcctggtcgtggctaaggtggaaaaaggaaagtctaagaagctcaagagcgtgaaggaactgctgggtatcaccattatggagcgcagctccttcgagaagaacccaattgactttctcgaagccaaaggttacaaggaagtcaagaaggaccttatcatcaagctcccaaagtatagcctgttcgaactggagaatgggcggaagcggatgctcgcctccgctggcgaacttcagaagggtaatgagctggctctcccctccaagtacgtgaatttcctctaccttgcaagccattacgagaagctgaaggggagccccgaggacaacgagcaaaagcaactgtttgtggagcagcataagcattatctggacgagatcattgagcagatttccgagttttctaaacgcgtcattctcgctgatgccaacctcgataaagtccttagcgcatacaataagcacagagacaaaccaattcgggagcaggctgagaatatcatccacctgttcaccctcaccaatcttggtgcccctgccgcattcaagtacttcgacaccaccatcgaccggaaacgctatacctccaccaaagaagtgctggacgccaccctcatccaccagagcatcaccggactttacgaaactcggattgacctctcacagctcggaggggatgagggagctcccaagaaaaagcgcaaggtaatggccacgaaaaatattcttcaaaatataaatgcacaatggaaggagaatgacaacagatcaccaagtagaaagcgacggttagatgacgtcactgaagaatcacaattaccttccacgaccaaacgacgtcatcttcaaacaaatacaaacgttgtaaattccacaggattgaaacaaggcctgacaaatgtgaaaaattcaataaatccaaagaacaaatcaataaaaaatttcttttctgatattccacgtgtgtcatgtactaaatctgaaaagattcaaatttttaaagaagctaagaaaactccaaaaaagaatgcaaccactcagacaaggagtgaagctgaagaattggtctgcagtgatcaacccagtgaaaaatattgggaactcttagccgaggagcgaaggaaagggttgtaa

>Noggin

ATGAATTTCGCTACTTGTTTTGCTACTTTAATGACGTGGATTGCCATCTCGGGATCGGGAATTTTATTCGTTGTCGGCCAGCATTACAAGCACCTACGCCCGCAGCCACATGATAAACTTCCTTTGCAAGATATCCCAGAAAATCCGAACCCGCTCTACAACCCCCGGCCCACTGACATCGATGTACGAAAGCTACGAGCAAAGTTAGGAGTCGATTACCTGCCTAACTTCATGTCGCCGCGAGACCCAGGAATCGTGGCCGTACCCACCGTCCATGTTAAGCAAAAGCCTCGACCCACAGCTGCGATCCGTCAAGTTTTCCGTCGTTTAACAAGGAATAAGTTTCCACGCCGGATGACCAGGCGAGTTATGAAGCACAAAAAGGTCATTCAATACTGGCTTTGGCAGGCGACATCTTGTCAGGTGCATTACCGATGGAAAGACCTTGGAATCCGATATTGGCCAAGGTTCATAAAAGAGGGGTACTGCGACACGACCCAAAGTTGTAGTATACCTGCCGGGATGCGATGCCACCAGTCCGGCCAAACGAACCTTAAAATCCTGCGCTGGGTTTGCCAAGGAATCTCCGAGCAGAAGTATTGCTTATGGATTGAGATGCACTACCCGATTATCAGTGAGTGCCGTTGCCAATGCCCTTAA

>NK4

ATGATTCCTAGTCCGGTTGGATCGACTCCATTTTCGGTTAAAGACATTTTGAACCTGGAACGTCATCAAATGTCATGTCAAAGCAGCGCCGATGACCGAGACACGCCCCATGTGGTCAATGGCTATCCCGTGTCGTGTAATGACAGCATAAAAAGCGAAGATACTCAAGTGCTTGATATGCATTCGATATACCATCACAGTTCGATGGAGGCCACCCAGCAAGCATCAGGTGAATTAGTTAAAGGAAGATTCGAAAAGTTGAGCAAACCGCACTCAAGACAAATCGAAACAAACAATATGGCTGCCTCCGATTCACAGTTCGACATGTTTTCGCAAAATGAAAACGCAAAATATTCTAAGGGGGCAATAGACACAAGGTATGCAGAAGCAACAAACACAGCAGACACGGCAGCGTCGAACCTTTTGCATTTTTCTCAAAATTTTCACCAAATTCCGTACGGGGAAAGTAGGGCACTGGTGGCGCCATTCGACATGCCTCGTTACGGTGGGGAGTATTTCAGTCCGACAAGCAACTACAGTGGAGGTTACACTCACCCCTATGACAACCCACCTAGCAATGTTGAAGCCTCACGATCGTCGTCGTTTCATTCACAGCAGTTTACGAAACCAAACGACGACGTTATTAGTACAACACACTCAACGTCCTCTGCCACAGCAGGAAACTTTTACCACAGACCGGCGAACAACTTCGAGCCTGTGAGAGAGCCAGGGCTGGAGAGATGTCCTGACAACGGAGACGAATTTTACACCAAAACTGAATCTCCGTTTCCCGCATTCAACCAACAGCATCAACCCACACCACAAAGCGAGCATCACAGCATGCCCAGCTACGACGAAATGAACGAGAAGATGTACACCGACCAGACATCCGAAACCGATATAACCAATAACTCAATGTTCGATTCGTCAAGTTCGGAGGCGGTGGTCGTCGATTTTCCATCACCATCCGGATCGCCAAACAAACAACATGAAGACGGAAGATTCGGAGGATCCTGTACGGATGAATCACTTCCGAGGACTTTCTCCGCCGCCGACATCCGGCAGGATGAATCAGTTACTCAAGTCACGGATTCGGAAAAGTCGGACAACGTCAGCACATGTAGCACTAGCCCCGAAGAAACTGAAAAAAAGAAAACAGAAGATGACACGCTAAAATCCCGACACCGGACCAGGCGAAAACCGCGTGTGCTGTTCTCCCAGGCACAAGTATTTGAGCTTGAGCGAAGATTTAAACAACAAAGATACTTGTCTGCTCCGGAAAGAGAACATCTGGCCCAGATTTTAAAACTGACTTCCACCCAGGTCAAGATCTGGTTCCAAAATCGTCGATACAAATGTAAACGAATGCGACAAGACAAAACCTTAGAGCTGGCCAGTATTGGACCTCCACGAAGAGTTGCAGTTCCTGTTTTAGTTCGCGACGGGAAGTCATGTCTCGGTGGGCCTGGTCAAGTATCACATAACGCGGCTCCATACAATGTGACTGTTACACGCTACCACAATTACCCAAGTTCATATAGCTCTTGCGGTTATAACTCCTACCCCCCTAACCCACCCGGTTACGGGCCAAGCGCTGCCGCAGCAGCAGCTGTAGCAGGAGCGGCTGCTGCAGCATACGGGAACGCTGCTACGAACGGTTATAACGGATTTCCGCCGAGCATGAACGGTGGACAGCCCGGTGCGAGCCCCTATCCTACTCCAGGGCTGCACCAACAGCACCCACCCTCTATGGGGGTCGGGAACTCACCGTTTCAGACCTCGACTGTACCTACAGGCCACCATATGTCTCAAGGGGGGTTGCACCATCACACACCAACGGACTATATGTATAAGCTTGGGCTGTGCACGACTAGTTGA

>NK4::VP16 EcoRI VP16 domain

ATGATTCCTAGTCCGGTTGGATCGACTCCATTTTCGGTTAAAGACATTTTGAACCTGGAACGTCATCAAATGTCATGTCAAAGCAGCGCCGATGACCGAGACACGCCCCATGTGGTCAATGGCTATCCCGTGTCGTGTAATGACAGCATAAAAAGCGAAGATACTCAAGTGCTTGATATGCATTCGATATACCATCACAGTTCGATGGAGGCCACCCAGCAAGCATCAGGTGAATTAGTTAAAGGAAGATTCGAAAAGTTGAGCAAACCGCACTCAAGACAAATCGAAACAAACAATATGGCTGCCTCCGATTCACAGTTCGACATGTTTTCGCAAAATGAAAACGCAAAATATTCTAAGGGGGCAATAGACACAAGGTATGCAGAAGCAACAAACACAGCAGACACGGCAGCGTCGAACCTTTTGCATTTTTCTCAAAATTTTCACCAAATTCCGTACGGGGAAAGTAGGGCACTGGTGGCGCCATTCGACATGCCTCGTTACGGTGGGGAGTATTTCAGTCCGACAAGCAACTACAGTGGAGGTTACACTCACCCCTATGACAACCCACCTAGCAATGTTGAAGCCTCACGATCGTCGTCGTTTCATTCACAGCAGTTTACGAAACCAAACGACGACGTTATTAGTACAACACACTCAACGTCCTCTGCCACAGCAGGAAACTTTTACCACAGACCGGCGAACAACTTCGAGCCTGTGAGAGAGCCAGGGCTGGAGAGATGTCCTGACAACGGAGACGAATTTTACACCAAAACTGAATCTCCGTTTCCCGCATTCAACCAACAGCATCAACCCACACCACAAAGCGAGCATCACAGCATGCCCAGCTACGACGAAATGAACGAGAAGATGTACACCGACCAGACATCCGAAACCGATATAACCAATAACTCAATGTTCGATTCGTCAAGTTCGGAGGCGGTGGTCGTCGATTTTCCATCACCATCCGGATCGCCAAACAAACAACATGAAGACGGAAGATTCGGAGGATCCTGTACGGATGAATCACTTCCGAGGACTTTCTCCGCCGCCGACATCCGGCAGGATGAATCAGTTACTCAAGTCACGGATTCGGAAAAGTCGGACAACGTCAGCACATGTAGCACTAGCCCCGAAGAAACTGAAAAAAAGAAAACAGAAGATGACACGCTAAAATCCCGACACCGGACCAGGCGAAAACCGCGTGTGCTGTTCTCCCAGGCACAAGTATTTGAGCTTGAGCGAAGATTTAAACAACAAAGATACTTGTCTGCTCCGGAAAGAGAACATCTGGCCCAGATTTTAAAACTGACTTCCACACAAGTCAAGATCTGGTTCCAAAATCGTCGATACAAATGTAAACGAATGCGACAAGACAAAACCTTAGAGCTGGCCAGTGAATTCGCACCACCGACCGATGTCAGCCTGGGGGACGAGCTCCACTTAGACGGCGAGGACGTGGCGATGGCGCATGCCGACGCGCTAGACGATTTCGATCTGGACATGTTGGGGGACGGGGATTCCCCGGGTCCGGGATTTACCCCCCACGACTCCGCCCCCTACGGCGCTCTGGATATGGCCGACTTCGAGTTTGAGCAGATGTTTACCGATGCCCTTGGAATTGACGAGTACGGTGGGTGA

>NK4::WRPW WRPW domain

ATGATTCCTAGTCCGGTTGGATCGACTCCATTTTCGGTTAAAGACATTTTGAACCTGGAACGTCATCAAATGTCATGTCAAAGCAGCGCCGATGACCGAGACACGCCCCATGTGGTCAATGGCTATCCCGTGTCGTGTAATGACAGCATAAAAAGCGAAGATACTCAAGTGCTTGATATGCATTCGATATACCATCACAGTTCGATGGAGGCCACCCAGCAAGCATCAGGTGAATTAGTTAAAGGAAGATTCGAAAAGTTGAGCAAACCGCACTCAAGACAAATCGAAACAAACAATATGGCTGCCTCCGATTCACAGTTCGACATGTTTTCGCAAAATGAAAACGCAAAATATTCTAAGGGGGCAATAGACACAAGGTATGCAGAAGCAACAAACACAGCAGACACGGCAGCGTCGAACCTTTTGCATTTTTCTCAAAATTTTCACCAAATTCCGTACGGGGAAAGTAGGGCACTGGTGGCGCCATTCGACATGCCTCGTTACGGTGGGGAGTATTTCAGTCCGACAAGCAACTACAGTGGAGGTTACACTCACCCCTATGACAACCCACCTAGCAATGTTGAAGCCTCACGATCGTCGTCGTTTCATTCACAGCAGTTTACGAAACCAAACGACGACGTTATTAGTACAACACACTCAACGTCCTCTGCCACAGCAGGAAACTTTTACCACAGACCGGCGAACAACTTCGAGCCTGTGAGAGAGCCAGGGCTGGAGAGATGTCCTGACAACGGAGACGAATTTTACACCAAAACTGAATCTCCGTTTCCCGCATTCAACCAACAGCATCAACCCACACCACAAAGCGAGCATCACAGCATGCCCAGCTACGACGAAATGAACGAGAAGATGTACACCGACCAGACATCCGAAACCGATATAACCAATAACTCAATGTTCGATTCGTCAAGTTCGGAGGCGGTGGTCGTCGATTTTCCATCACCATCCGGATCGCCAAACAAACAACATGAAGACGGAAGATTCGGAGGATCCTGTACGGATGAATCACTTCCGAGGACTTTCTCCGCCGCCGACATCCGGCAGGATGAATCAGTTACTCAAGTCACGGATTCGGAAAAGTCGGACAACGTCAGCACATGTAGCACTAGCCCCGAAGAAACTGAAAAAAAGAAAACAGAAGATGACACGCTAAAATCCCGACACCGGACCAGGCGAAAACCGCGTGTGCTGTTCTCCCAGGCACAAGTATTTGAGCTTGAGCGAAGATTTAAACAACAAAGATACTTGTCTGCTCCGGAAAGAGAACATCTGGCCCAGATTTTAAAACTGACTTCCACACAAGTCAAGATCTGGTTCCAAAATCGTCGATACAAATGTAAACGAATGCGACAAGACAAAACCTTAGAGCTGGCCAGTCAGATCAAGGAGGAGGAGCAGCCCTGGCGGCCCTGGTAATAATGA

>Msx

ATGACAGTAAACGAATCCGATCGCATTCAATCCTCTGTATTGTCGGAATGTGTATCCGAACCTGAAGTGACGTCATCGACAGTGACGTCACCAAAACCTAAGGAAGAGTCAATTAACCACGACAAAGATTCGAGTTTTAATAATTCAGAAACACAAGAAAATCCACCGAAAAAAATTAAAAAGGAAAAACTTAATTTTAGCATTGAgTTCCTACTTTCCAAACCTGAAAGAAAAACCCGGACACCTCCAATCCAATCCACAGATGCTTCCCTTCAACACAAACCTTATTTTTCCGGATATTTCACAAGTTACAACCCAACTTTTGACCCTTTTTCTATGTGGGGTAAATCCCATTCACCCCCCAAGCCCCAACTCGATTACCCAGCTATAACGTCAAAGGAATCGTCACCCACgcgtTCTGACGTCACAGAAACTGATGATTGTAGTTACGAGTCGACGTCGCCGAAGTCTAATGACGTCATTCCGACGTCACCGAATGCTTTCCGTATATCTAAATGTTCACTTCGTAAACACAAACCAAACCGAAAACCCCGCACACCGTTCAGCACCGAGCAACTACTTTCATTGGAACGAAGATTTCAGGACAAACAATATCTTTCTATAGCAGAACGTGCAGAgTTCTCTGCCTCTCTTGCATTATCTGAAACTCAGGTGAAAATTTGGTTTCAAAACCGTCGCGCTAAAGCAAAACGTCTTCACGAAGCTGAGTTTGAAAAAGTGAAGCTGGCTGCTGCAGCAGCAGCATATACTGACTTATTACAACCGCCCACAAAACCTCACGCCTTATACCCCGGTTACGGTGTAAGAGGAATACCCACAGACCCAGCAATGCGATTGGGGCATACTGGGCCACTAGCCCGAGGTGTAACAGCATCGCCGAGCCTAAACGCATTTTCAAACTTTGCCCAGAGTCGTTGTCATCGACCAGCGACTATCTCCTACTACCCAACTGAGAGTCGATAG

>Chordin

ATGATGATGACGTCATGCTATTACGTCACGGTGGTATTCATTGCTATGACGTCAGGTATATGTTCTGGCTTTCTGCAGATACATCCAGCAGAAGAAGCATTGCCGTCTAATAACCCGAGAGGTTGTGTATTTGGTCGTCAGTTCCACGTCATCGGATCATCATGGCACCCAAACCTTGGTCGTCCATTTGGAATCATGTATTGTGTCACGTGCCAGTGCGTCAAAGAGACCACGGGACGTTTCAAACCAGACGGACAACCAAAGGAAGTTACTCGGGTCACGTGTCATGACGTAAGAAAGGACTGCCCCCACGTCACGTGCAAAGGGGCCAAGGTTCCAAGGGGAGGATGCTGTAAAGTTTGCCCAGACGATACAGATACAATTGTGACGTCAGCCGAAGAATTTTCATCGAAAAATGGTGATGTTCCGCTGCCATTGGATATCACTGAAGAAAAATCAATTAACCACGACGACTCAAAGGCTGGTGTTGATGAGAAGACAAAAAGTTATGTTGCACTGTTAACCTCAAACGATGGTGGGCCATTGTTACGAGCTACGTTTAACCTGCACAGGGATAATCTGCATTTTACGATCCAGTATTTGAGGTCAACCAAACCCACCATTATAGAAATGCTGACTAATGATGACGTCACAATATACACTCATGTGACTGAAATGTCACCCAAGCCAGGACAACCTATTGTTTGCGGGGTATGGCGTAATCTACCAAAACACGTTGTTAGCTCTTTAGATAAAAGGTTGTTGACTCTCAGAGCTGTTTTTGTCCAAACTGGTGCCAATATGACATCACAAGGTCAAATAATACGACACAGAGCATTGAGTCGAGAAACTTTTAGTTCGATTTTATCCACGACACCGCTAGATGGAGCTAGTGGAGCAATTGCGATGATCACCCTTGGCCGACATCGATTCAATAAACTTCATTTTGCCATTTTACTCAGACGATCTTCTGTGACATCACAACCCACCACAATCACTGTGCGATTACTCAACAATCAAAACAGAACGTTGCGATCAACCGAGAAAACTCTTCAACCCTCAAGTAATGAGCTTGCTGATGTGTGGTCGACAAATTCTGAAATCCTTCGAAGATTGGGTGATGGTTCACTTCATATACAGATGGTTGCTAGGACAACAGGGTCACATGATAAACAAAGCTACGTCGGCAGCGTGAGAGCAAAAACAACTTGTGACCTGTTGCAAGCTGTTTTATCTGGTAGTGACTGTAGCCTACCAGCCAGTACAGGAGCAGCTGGATCCGCGATTCTGAATATTAATTCTGACAACACTGTTTCTTACAAGGTGGCGCTAGTGGGTATTTCAAGTCGGGTGACCTCAATATCACTGCTTGGAGACTACAGAAAGAAGCGAACACGCGTAGTAGCCGGTCTCACCCCGCAATTTGTGAATGGAAAAGCACAAGGAAGACTGGAAAACCTTAATGGGAAAGAGATACATCTTTTACTAAGCAACCGTGTGCGTGTTGCCGTGGAAACTGAACAAAGCAATAGTGGAGAGCTGGGTGGTCATGTGACTTCTCTCCTTTATGGAGGCCACCAAGCTAGATATAAAGGCCTTCCAATTCCTCTTGCTGGTTCTCTAGTACGACCCCCTGTATTCACTGGAGCTGCAGGTCACGTTTGGCTTGAACTGAATGAGGATTGCCATCTTTATTATGAAATTGTTGTTTCTGGACTAAGCAAAGAGCGTGACACAACATTAGCCGCACATTTACACGGTCTCGCTGAAATAGCTGAAACAGAAACTGGACATAAACAACTTTTGAAAGGTTTCTATGGAAAAGAGGCGCGTGGAATATTAAGAGAATTATCCTCAGAAATATATGAACATCTCAACCAAGGCAATGCTTTTGTACAGATTGCAACAAAGTCAAACCCAGGTGGAGAAATACGTGGTCGTGTTCATATACCAAACACTTGCTCCAGGACAATAGTTAACAAGGCTAACCATAAACAAGATACTTCAGACACAGACCAGAATTCATTAAGTGTTCCAAGCTCTGATGTCACCACGTCACGGGTGACTGATCTCGATGCAGTGGTGCATGACGGGGTAATCCCAACATACACCGAGCGTCGTTTGGAGCTTGACCCATCCGCTTGTTACGTCATAGACGAATGGAAACCCCATGGTAGTGAATGGGCACCAGATTACGACCCAAAATGCACAATATGCGTTTGTGAGTCCGGTAGTTCACTTTGCGATCCTGTTTACTGCCCGCCACTAGAGTGCGCCAGACCAGTGGCCACCGAAGGGAGCTGTTGTCCAGTATGTAACGATGGGTTTGGGGGAATACCGACCAAACATGATGATGGGGCTTCTATAGGATCTCTGTTTGGGTCTCCATTTACACAACTGGATCAAAAAGAAGACGAAAAAGAAAAAGAGAAAGAAGGTTGCTACTACGGGGGCGATGGACAAATACACGCGATAGGTTCATGGTGGCATCCATATTTTAAACGATTCGGCCACGTATCGTGCGTGAACTGCACTTGTAGGCCAGACGGTGAAATTGTGTGTGGAAAAATTACGTGCCCGCCAGTAACGTGCTCAAATCCTATCAAACAAAATCCGAGAGATTGCTGTAAAACTTGCCCAGAGGTCACACGTCACGAACTACGTCACGAATCACGTGACCATCGCATGCAGGACGACAGCGTGTCCAGGATGTGTCAGTTTGGTCGTGAGCTACGAGCACCCGGGGAAAAATGGACGCTTGGATATCTGGGCAGAAGAAACCTTGAATGTATAGAATGCGAATGTGCTACCCATGGTGTGAAACACAAGTGCCGCAAAAACTGTCCAGCTGTCAGTCATGAATGCAGTCGAGTGGAACGAAACGTTTCCGGTTGCTGCCCATGGCGTTGCGCGGACCGTCAGAGGTCGACCGAGAGATTTCCGTTTTCGTAA

>Foxg>Foxg

(originally used as a rescue construct in Johnson et al. 2024)

Foxg -2863/-1 START Foxg CDS PAM/target mismatch disrupted EcoRI

gtttcattttccaacaaaacattataatgcgataagagaattggccatctttaccacagaacttaaatttaacggaaatataaagttgcaacgtacgtaaataacagaccacccgtatttattgcgtgacattaatacattggatgtcaaagaacaaagaacttgttgtttacgtcatataggatccggtactagataaaccacttcaagaataagatttatgacatcataaccaaacatgagcgacgactttaattgggaaaaaatactttgaaatgttttttttatcagatatttgacaacaaaaatacaatagtcggaaaatattgtatttttcagttttatttgttcatatttaatatataagcacacctaacacctggctgataatgggacgtcactgggttaaaatagaaatcatataaagcacaagttcaattggtttcgggctaaaaattaaatcctttttcggtaaaccgaccgtttgtcacatcaaagccacagcgagcagtgataagagccgagtgtttttatgtaaataggtttaggtaactcgtgtctgcttgatttggttgtaagattaattactcacagtgttacacatgttcccacaaacccagagacattgtaactggggcgtagatgatacgttataaaatataaaccaacacatgttgtataatataagtgcactcattttatccgcagagtgttataacgattttcgcttttaaaatatatagtagggtgggggaagatgggacacctttagcacataatatccaaatgttctcatcgcgttatgaaccatttcaacggtctattgtcgtaaggatacggttttgtaattctttgaatgttttttgtttactaccaagtgggacgagaaattagaatgaaaaggtgtcccatctttccccgctctaatatataacttttttaatctgtgtaatgtttcatttaaacatgttctgggctattgagttcttaaaacttatcaaacgtatgttggtatatctgcccggacctttattattaaacaccaaaaccacatagaaaaagtaactattaaacgaaattctgtctatgtcagccgttatactgacttgcagtgtgtatagatatcgttaggctctatttggtggtaaacaaagaatatttacagaattatataaccgtattcttacgactgtaaaagagtgttgttaattgtttaaaacacgaataaaaaatatgggagatcgtgtgctgatggttttcaccttatcacagttcagataaaaaattaaaagttctcaaaacaaactataggaacaaatatgtttttgatttatattattgcaaaaatactatgttaaattggcctacctatttccaattaaaaaaagtcacattttaattaattttacggaaacactgtacgttgtcagtgattttagacctgatcttataattgagtctggagtaaaacagaccttgtgattgacagtttgaattataagtatcagaaaacattgatcgtagatgtaaccaaagctctcatcagttgtggaccaaagtaaccacaaaacctacctaatttttgcagtgttgcctttagactaaacttttatttcagtgttgccatttaaaaaaaaagctagatgtgaggtcggtgaataaaaagggtaaattattagttatatgagcgtagtttaagctataactagttaatacagatatattaaatcatgcagtttgcgtacaaattatattaattaaagtccattaaaaatgtattcaatattacaaaaaaatcaatttctattaagaaaaacattaaaaaattggaattttttgacattttaatcgtggcaacatgacaaacttatgttagcagattatcacgccatcttagcatacagcacgtgtgaagcaagatcaaataaccaaggtgtcaagtgaaagttacagggcacaattatttaaccaagctaacgtaaatgaaaaacccaggcaaccggcttattactgctggctacttggtcagtgtggttaacatatatcaacggaaagccccgctctatatcatgacgtattaactagtattaacaagcccaacattgtacggtgtcgtgtaaaaccttatatggcatggagcttgttttgcggaaagctattttgttttttaatctgaaaatatgagattgagtgtggtatgggaaaatgtactcctatagacacacctggtgtattcacacaccttatcctcactatgttgagttatagcatgggtaattcttatccaccacacatggtgtatttactgtattcatacttagttataacagtattgtatatttaaatattgccaaaatccgtcgtagagtgacaaaaacaatcttgaaatactgcaaagtcattaggcgtattatctcacactctaaatcacttaaacacaataaagtcgcgagttttaaaagcctagatctaataataattgcaaatcaaacgcattatgtccattagttgtattttctttgtcgatttagtatccgggttatccttgttaacaaggtgtttgtgtgtctaagtatgtccatgattacgtcttcctaccagaggagtcgtgttgttttagtcaatacggtggtaattatccgattattgtgatatagcgagttacaggcgccctcagctcacggtggaagtgctaacttgtaagtcttaatgcatttctcttttcttcaaaggtttttgagtagattttcgtttgtaattataccggcgtctccttacgctgtaatccacgcttttaattgtgacgtggattgacagaaagagtagcgcgagagagaaacgcggcatacagaatATGACGAACGACGCGGCCGAGTCTGGTCATTCAAAGCGAGAAAACTTCATAGAGATGTCGCCCGAGTATTCAACGCTGATCGCCGCCGAAGATCGAAGCTCAGTTCCCagacgcGAaGACGCGCAAGTGCCGAGGTTTGAGAATATACAAAATGGTGGTGCCGATATAGAAAACGGTGAAGTGTCTCCAATACAAAAAGATATTGCAAACCAAGAGCTCAATGACGTAGCAATTAATATGACGTCACCTCAGAAACAAACAAACAACGACAATATAGAAGAAAAATGTCCAAAAGACCAGAAACCGTCAACAAGTCCACCGAGTAACAAGTACGGTAAGAAACCGCCATATTCATACAACGCCTTGATAATGATGGCCATCAAAAAGAGCCCACGAAAGCGACTTACACTAAGTCAGATCTACCAATACATAACAACTACATTCCCGTACTACAAAGAAAATAAACAGGCGTGGCAGAAcTCTATCAGACACAACTTATCGTTGAACAAATGCTTTGTGAAAGTACCGAGGCACTACGACGACCCCGGGAAGGGTAACTATTGGATGTTAGACCCGTCTAGTGATGATGTATACATTGGTAGTAGCACCGGTAAACTGAGACGGAGAAGTTCAAGCAGTCAAGCAAGGGGTCGTTTAGCATTACGACGAAGAACATTCGCCCAAGTATTTGGATCGCCGCAAGATATTTTACAACACGACCCCCCACAACAGATAATAAGAACAGACGTGACGTCACGAGCGGCCATGTTACGTCATCATGACGCAGCACGAACGATGCTGGGTTCTGGAATCCCACAGCGGGCAGATCCCTACCCTATGTTCCAACACACGCGGTTCCCTATGACTACAGAAACTGATTCAAGATACAGACAATTGTATAGAGCGAGATTGGAACAATACTACGCGCATCTCGCGTCCTCTGCACTGTTCGGGCACATGCAAGCTACTGCATTAAACGCCCAGCCACGTATTGTTAAAAGCAACCCCGCCTCTCCTGAGCCTGTGCATTCCCATTACGAACAAACTACGTCACCAAACACCGCACCTCCaCGaTCTGACACCTCCACACCACCGCGAGGGATCGAACGACCGTGGGCATCACCGCCGCGTAGGAGGTGTGACGTCACAAAGCATGAAACAATTGACCGCGACGTCGTGTCGCGATGCTCCTCCGAATCTTCTAATGAATCATCAAGAAACAAGAGAGAAGTCACCGAAAACTCTACGAAcTCACCGATACAGCAACTTAGTTCAAGAGGAGGGTTACCGTTCTACCTTACACCAACCCCGAACCCTTGCCCTAATTTCTTAATGCCCAACACTGCCGGGTTAGTTCCAAATCCAAGTTACCCCTTCTTTTTTCCCCAACCATTCCACCCAGCCTTGGCGTTTCTCTCTCGGCCACAAACTGTTGCGTCATCATCGCAATTGTGA

**Protein coding sequences for sequence alignment**

>NK4 (Ciona robusta)

MIPSPVGSTPFSVKDILNLERHQMSCQSSADDRDTPHVVNGYPVSCNDSIKSEDTQVLDMHSIYHHSSMEATQQASGELVKGRFEKLSKPHSRQIETNNMAASDSQFDMFSQNENAKYSKGAIDTRYAEATNTADTAASNLLHFSQNFHQIPYGESRALVAPFDMPRYGGEYFSPTSNYSGGYTHPYDNPPSNVEASRSSSFHSQQFTKPNDDVLSTTHSTSSATAGNFYHRPANNFEPVREPGLERCPDNGDEFYTKTESPFPAFNQQHQPTPQSEHHSMPSYDEMNEKMYTDQTSETDITNNSMFDSSSSEAVVVDFPSPSGSPNKQHEDGRFGGSCTDESLPRTFSAADIRQDESVTQVTDSEKSDNVSTCSTSPEETEKKKTEDDTLKSRHRTRRKPRVLFSQAQVFELERRFKQQRYLSAPEREHLAQILKLTSTQVKIWFQNRRYKCKRMRQDKTLELASIGPPRRVAVPVLVRDGKSCLGGPGQVSHNAAPYNVTVTRYHNYPSSYSSCGYNSYPPNPPGYGPSAAAAAAVAGAAAAAYGNAATNGYNGFPPSMNGGQPGASPYPTPGLHQQHPPSMGVGNSPFQTSTVPTGHHMSQGGLHHHTPTDYMYKLGLCT

>NKX2-5 (Homo sapiens)

MFPSPALTPTPFSVKDILNLEQQQRSLAAAGELSARLEATLAPSSCMLAAFKPEAYAGPEAAAPGLPELRAELGRAPSPAKCASAFPAAPAFYPRAYSDPDPAKDPRAEKKELCALQKAVELEKTEADNAERPRARRRRKPRVLFSQAQVYELERRFKQQRYLSAPERDQLASVLKLTSTQVKIWFQNRRYKCKRQRQDQTLELVGLPPPPPPPARRIAVPVLVRDGKPCLGDSAPYAPAYGVGLNPYGYNAYPAYPGYGGAACSPGYSCTAAYPAGPSPAQPATAAANNNFVNFGVGDLNAVQSPGIPQSNSGVSTLHGIRAW

>Msx (Ciona robusta)

MTVNESDRIQSSVLSECVSEPEVTSSTVTSPKPKEESINHDKDSSFNNSETQENPPKKIKKEKLNFSIEFLLSKPERKTRTPPIQSTDASLQHKPYFSGYFTSYNPTFDPFSMWGKSHSPPKPQLDYPAMTSKESSPTRSDVTETDDCSYESTSPKSNDVIPTSPNAFRISKCSLRKHKPNRKPRTPFSTEQLLSLERRFQDKQYLSIAERAEFSASLALSETQVKIWFQNRRAKAKRLHEAEFEKVKLAAAAAAYTDLLQPPTKPHALYPGYGVRGLPTDPAMRLGHTGPLARGVTASPSLNAFSNFAQSRCHRPATISYYPTESR

**Previously published sgRNAs:**

NK4.3 (targets exon 2, from Gandhi et al. 2017)

**GAACCAGATCTTGACCTGGG (G+N19)**

Control (targets no *Ciona* sequence, from Stolfi et al. 2014)

**GCTTTGCTACGATCTACATT (G+N19)**

Defcab.47

(used as control, previously Efcab6-r.47 from Gibboney et al. 2020)

**GGCCAGCCAAGCCAGCAGGG (G+N19)**

**Newly designed sgRNAs:**

NK4.113 (targeting exon 1)

**GGACCGAGACACGCCCCATG (G+N19)**

Chordin.exon3.4

**GTCTGGTTTGAAACGCCCAG (G+N19)**

Chordin.exon3.116

**GAAGGGGCCAAGGTACCCAG (G+N19)**

Chordin.exon5.29

**GCACTGTTAACCAGCAATGA (G+N19)**

*PCR primers for amplicon sequencing by NGS*

| **Gene + exon** | **Forward primer** | **Reverse primer** |
| --- | --- | --- |
| NK4 exon 1 | ACTGCTGAAACGCTATTTGTAC | GAGGCAGCCATATTGTTTGT |
| Chordin exon 3 | ACGAGCCGTATTACGTCAC | TCTTGACGGTGTTGTAGCC |
| Chordin exon 5 | CATCCATGCTTTTGGGTAATG | AGCAACAACAGGAGCATTG |

**Electroporation mixes for perturbation experiments**

**(per 700 ul of total cuvette volume)**

NK4 CRISPR to look at ACCs (CryBG reporter):

35 ug Foxc>Cas9::Geminin N-terminus

10 ug Foxc>H2B::mCherry

50 ug U6>NK4.3 sgRNA

35 ug CryBG>Unc-76::GFP

Negative control for the NK4 CRISPR/ACC experiment above:

35 ug Foxc>Cas9::Geminin N-terminus

10 ug Foxc>H2B::mCherry

50 ug U6>Control sgRNA

35 ug CryBG>Unc-76::GFP

NK4 CRISPR to score “Cyrano” phenotype:

35 ug Foxc>Cas9::Geminin N-terminus

10 ug Foxc>H2B::mCherry

50 ug U6>NK4.3 sgRNA

50 ug Islet int1a + bpFOG>Unc-76::GFP

Negative control for the NK4 CRISPR phenotype experiment above:

10 ug Foxc>H2B::mCherry

50 ug Islet int1a + bpFOG>Unc-76::GFP

NK4 CRISPR to look at ICs (C11.360 reporter):

35 ug Foxc>Cas9::Geminin N-terminus

10 ug Foxc>H2B::mCherry

50 ug U6>NK4.3 sgRNA

50 ug CryBG>Unc-76::GFP

Negative control for the NK4 CRISPR/IC experiment above:

35 ug Foxc>Cas9::Geminin N-terminus

10 ug Foxc>H2B::mCherry

50 ug U6>Control sgRNA

50 ug CryBG>Unc-76::GFP

NK4 CRISPR to look at OCs (L141.36 reporter):

35 ug Foxc>Cas9::Geminin N-terminus

10 ug Foxc>H2B::mCherry

50 ug U6>NK4.3 sgRNA

50 ug L141.36>Unc-76::GFP

Negative control for the NK4 CRISPR/OC experiment above:

35 ug Foxc>Cas9::Geminin N-terminus

10 ug Foxc>H2B::mCherry

50 ug U6>Control sgRNA

50 ug L141.36>Unc-76::GFP

NK4 CRISPR to look at PNs (Seldom reporter):

35 ug Foxc>Cas9::Geminin N-terminus

10 ug Foxc>H2B::mCherry

50 ug U6>NK4.3 sgRNA

50 ug Seldom>Unc-76::GFP

Negative control for the NK4 CRISPR/PN experiment above:

35 ug Foxc>Cas9::Geminin N-terminus

10 ug Foxc>H2B::mCherry

50 ug U6>Control sgRNA

50 ug Seldom>Unc-76::GFP

NK4 CRISPR to look at metamorphosis at 44 hpf:

35 ug Foxc>Cas9::Geminin N-terminus

10 ug Foxc>H2B::mCherry

50 ug U6>NK4.3 sgRNA

50 ug Seldom>Unc-76::GFP

Negative control for the NK4 CRISPR/44 hpf experiment above:

35 ug Foxc>Cas9::Geminin N-terminus

10 ug Foxc>H2B::mCherry

50 ug U6>Control sgRNA

50 ug Seldom>Unc-76::GFP

Noggin overexpression looking at ACCs (Seldom reporter):

70 ug Foxc>Noggin

10 ug Foxc>H2B::mCherry

35 ug CryBG>Unc-76::GFP

Noggin overexpression looking at ICs (C11.360 reporter):

70 ug Foxc>Noggin

10 ug Foxc>H2B::mCherry

50 ug C11.360>Unc-76::GFP

Noggin overexpression looking at OCs (L141.36 reporter):

70 ug Foxc>Noggin

10 ug Foxc>H2B::mCherry

50 ug L141.36>Unc-76::GFP

Noggin overexpression looking at PNs (Seldom reporter):

70 ug Foxc>Noggin

10 ug Foxc>H2B::mCherry

50 ug Seldom>Unc-76::GFP

Noggin overexpression to quantify NK4 reporter activity:

70 ug Foxc>Noggin

70 ug Islet int1a + -473/+9>mCherry

60 ug NK4 int1.1 + bpFOG>Unc-76::GFP

Negative control to compare to Noggin quantification above:

70 ug Foxc>lacZ

70 ug Islet int1a + -473/+9>mCherry

60 ug NK4 int1.1 + bpFOG>Unc-76::GFP

Noggin overexpression for Msx reporter imaging:

70 ug Foxc>Noggin

10 ug Foxc>H2B::mCherry

70 ug Msx>CD4::GFP

NK4 CRISPR for Msx reporter imaging:

35 ug Foxc>Cas9::Geminin N-terminus

10 ug Foxc>H2B::mCherry

70 ug Msx>CD4::GFP

50 ug U6>NK4.3 sgRNA

Negative control for Nk4 CRISPR/Noggin overexpression above:

35 ug Foxc>Cas9::Geminin N-terminus

10 ug Foxc>H2B::mCherry

70 ug Msx>CD4::GFP

50 ug U6>Control sgRNA

NK4 overexpression looking at papilla flatness and Islet reporter:

50 ug Foxc>NK4

10 ug Foxc>H2B::mCherry

50 ug Islet int1a + bpFOG>Unc-76::GFP

Negative control for flatness/Islet reporter experiment above:

70 ug Foxc>lacZ

10 ug Foxc>H2B::mCherry

50 ug Islet int1a + bpFOG>Unc-76::GFP

Foxc>Noggin alone:

70 ug Foxc>Noggin

10 ug Foxc>H2B::mCherry

50 ug Islet int1a + bpFOG>Unc-76::GFP

Foxc>Noggin + Foxc>NK4 to compare to Noggin alone above:

70 ug Foxc>Noggin

70 ug Foxc>NK4

10 ug Foxc>H2B::mCherry

50 ug Islet int1a + bpFOG>Unc-76::GFP

NK4 CRISPR for Foxg in situ hybridization:

35 ug Foxc>Cas9::Geminin N-terminus

10 ug Foxc>H2B::mCherry

50 ug U6>NK4.3 sgRNA

Negative control for Foxg in situ hybridization:

35 ug Foxc>Cas9::Geminin N-terminus

10 ug Foxc>H2B::mCherry

50 ug U6>Control sgRNA

NK4 CRISPR for Islet reporter at 11 hpf:

35 ug Foxc>Cas9

50 ug U6>NK4.3 sgRNA

70 ug Islet int1a + -473/+9>Unc-76::GFP

Negative control to compare at 11 hpf above:

35 ug Foxc>Cas9

50 ug U6>Defcab.47 sgRNA

70 ug Islet int1a + -473/+9>Unc-76::GFP

NK4 CRISPR to look at Foxg>GFP in larva:

35 ug Foxc>Cas9::Geminin N-terminus

10 ug Foxc>H2B::mCherry

50 ug U6>NK4.3 sgRNA

70 ug Foxg>Unc-76::GFP

NK4 overexpression to look at Foxg>GFP:

50 ug Foxc>NK4

70 ug Foxg>Unc-76::GFP

10 ug Foxc>H2B::mCherry

Negative control to compare to NK4 experiment above (Foxg>GFP):

70 ug Foxg>Unc-76::GFP

10 ug Foxc>H2B::mCherry

Foxg>Foxg looking at ACCs (originally as a CRISPR rescue experiment):

50 Foxg>Foxg

40 ug Foxc>Cas9

25 ug U6>Foxg.1.116 sgRNA

25 ug U6>Foxg.5.419 sgRNA

10 ug Foxc>H2B::mCherry

35 ug CryBG>Unc-76::GFP

Foxg>Foxg looking at ICs:

50 ug Foxg>Foxg

10 ug Foxc>H2B::mCherry

50 ug C11.360>Unc-76::GFP

Foxg>Foxg looking at OCs:

50 ug Foxg>Foxg

10 ug Foxc>H2B::mCherry

50 ug L141.36>Unc-76::GFP

Foxg>Foxg looking at PNs:

50 ug Foxg>Foxg

10 ug Foxc>H2B::mCherry

50 ug Seldom>Unc-76::GFP

Foxg>Foxg + Foxc>NK4 to see if flat or Cyrano:

10 ug Foxc>H2B::mCherry

50 ug Foxg>Foxg

50 ug Foxc>NK4

50 ug Islet int1a + bpFOG>Unc-76::GFP

NK4::VP16 overexpression to look at Foxg reporter and papilla:

70 ug Foxg>Unc-76::GFP

10 ug Foxc>H2B::mCherry

50 ug Foxc>NK4::VP16

Negative control to compare to NK4::VP16 above:

70 ug Foxg>Unc-76::GFP

10 ug Foxc>H2B::mCherry

NK4::WRPW to look at flat papilla:

50 ug Foxc>NK4:WRPW

10 ug Foxc>H2B::mCherry

50 ug Islet int1a + bpFOG>Unc-76::GFP

Negative control to compare to NK4::WRPW above:

10 ug Foxc>H2B::mCherry

50 ug Islet int1a + bpFOG>Unc-76::GFP

NK4 and Msx co-overexpression:

50 ug Foxc>NK4

50 ug Foxc>Msx

70 ug Foxg>Unc-76::GFP

10 ug Foxc>H2B::mCherry

NK4 overexpression alone to compare to NK4+Msx above:

50 ug Foxc>NK4

70 ug Foxg>Unc-76::GFP

10 ug Foxc>H2B::mCherry

Chordin overexpression to look at papilla shape:

70 ug Foxc>Chordin

10 ug Foxc>H2B::mCherry

60 ug NK4 int1.1 + bpFOG>Unc-76::GFP

Negative control to compare to Chordin overexpression above:

70 ug Foxc>lacZ

10 ug Foxc>H2B::mCherry

60 ug NK4 int1.1 + bpFOG>Unc-76::GFP

Chordin CRISPR to quantify NK4 reporter expression:

35 ug Foxc>Cas9::Geminin N-terminus

10 ug Foxc>H2B::mCherry

50 ug NK4 int1.1 + bpFOG>Unc-76::GFP

25 ug U6>Chordin.exon3.4 sgRNA

25 ug U6>Chordin.exon3.116 sgRNA

25 ug U6>Chordin.exon5.29 sgRNA

Negative control to compare to Chordin CRISPR quantification above:

35 ug Foxc>Cas9::Geminin N-terminus

10 ug Foxc>H2B::mCherry

50 ug NK4 int1.1 + bpFOG>Unc-76::GFP

75 ug U6>Control sgRNA

Chordin CRISPR to look at dorsal papilla formation:

35 ug Foxc>Cas9::Geminin N-terminus

10 ug Foxc>H2B::GFP

30 ug Msx>H2B::mCherry

70 ug Foxg>Unc-76::GFP

25 ug U6>Chordin.exon3.4 sgRNA

25 ug U6>Chordin.exon3.116 sgRNA

25 ug U6>Chordin.exon5.29 sgRNA

Negative control to compare to Chordin CRISPR above:

35 ug Foxc>Cas9::Geminin N-terminus

10 ug Foxc>H2B::GFP

30 ug Msx>H2B::mCherry

70 ug Foxg>Unc-76::GFP

75 ug U6>Control sgRNA

NK4 CRISPR with NK4.113 sgRNA:

35 ug Foxc>Cas9::Geminin N-terminus

10 ug Foxc>H2B::mCherry

50 ug Islet int1a + -473/+9>Unc-76::GFP

50 ug U6>NK4.113 sgRNA

NK4 CRISPR to look at acetylated tubulin staining of PNs:

35 ug Foxc>Cas9::Geminin N-terminus

10 ug Foxc>H2B::mCherry

50 ug Seldom>Unc-76::GFP

50 ug U6>NK4.13 sgRNA

Negative control to compare acetylated tubulin staining to above:

35 ug Foxc>Cas9::Geminin N-terminus

10 ug Foxc>H2B::mCherry

50 ug Seldom>Unc-76::GFP

50 ug U6>Control sgRNA

*Ciona robusta* gene model ID table

| **Gene name and KH ID** | **KY21 ID** | **ANISEED gene ID** |
| --- | --- | --- |
| NK4 (KH.C8.482) | KY21.Chr8.898 | Cirobu.g00009121 |
| Seldom (KH.C4.78) | KY21.Chr4.267 | Cirobu.g00006902 |
| KH.C11.360 | KY21.Chr11.1038 | Cirobu.g00002187 |
| KH.L141.36 | KY21.Chr7.130 | Cirobu.g00011190 |
| KH.C14.116 | KY21.Chr14.663 | Cirobu.g00003524 |
| Msx (KH.C2.957) | KY21.Chr2.1031 | Cirobu.g00005203 |
| Chordin (KH.C6.145) | KY21.Chr6.382 | Cirobu.g00007664 |
| Noggin (KH.C12.562) | KY21.Chr12.737 | Cirobu.g00003129 |
| FOG (KH.C10.574) | KY21.Chr10.450 | Cirobu.g00001805 |
| Defcab (KH.C1.1218) | KY21.Chr1.2072 | Cirobu.g00000242 |
| CryBG (KH.S605.3) | KY21.Chr1.2268 | Cirobu.g00014792 |
| Islet (KH.L152.2) | KY21.Chr4.1164 | Cirobu.g00011396 |
| Foxc (KH.L57.25) | KY21.Chr12.158 | Cirobu.g00012813 |
| Foxg (KH.C8.774) | KY21.Chr8.693 | Cirobu.g00009441 |
